# 14-3-3 recruits keratin intermediate filaments to mechanically sensitive cell-cell contacts

**DOI:** 10.1101/349092

**Authors:** Richard A. Mariani, Shalaka Paranjpe, Radek Dobrowolski, Gregory F. Weber

## Abstract

Intermediate filament cytoskeletal networks simultaneously support mechanical integrity and influence signal transduction pathways. Marked remodeling of the keratin intermediate filament network accompanies collective cellular morphogenetic movements that occur during early embryonic development in the frog *Xenopus laevis*. While this reorganization of keratin is initiated by force transduction on cell-cell contacts mediated by C-cadherin, the mechanism by which keratin filament reorganization occurs remains poorly understood. In this work we demonstrate that 14-3-3 proteins regulate keratin reorganization dynamics in embryonic mesendoderm cells from *Xenopus* gastrula. 14-3-3 co-localizes with keratin filaments near cell-cell junctions in migrating mesendoderm. Co-immunoprecipitation, mass spectrometry and bioinformatic analyses indicate Keratin 19 is a target of 14-3-3 in the whole embryo and, more specifically, mesendoderm tissue. Inhibition of 14-3-3 results in both the decreased exchange of keratin subunits into filaments and blocks keratin filament recruitment toward cell-cell contacts. Synthetically coupling 14-3-3 to Keratin 19 through a unique fusion construct conversely induces the localization of this keratin population to the region of cell-cell contacts. Taken together, these findings indicate that 14-3-3 acts on keratin intermediate filaments and is involved in their reorganization to sites of cell adhesion.

## Introduction

Intermediate filaments (IFs) comprise a diverse group of structurally conserved proteins that assemble into long fibrils that function as a ‘cellular skeleton’. Found across many of the tissues of chordate eukaryotes, these cord-like macromolecules form intricate networks in both the cytoplasmic and nuclear compartments of cells. Keratins, a subtype of IF proteins, are expressed in epithelia as well as a few additional cell types, including cells of the early vertebrate embryo.

IFs have a well-established role in providing structural integrity within cells through binding to protein scaffolds that experience transmission of forces (Sanghvi-Shah and Weber, 2017). These interfaces include hemidesmosomes at cell-matrix contacts as well as desmosomes and classical cadherins at cell-cell junctions. IFs, including keratins, are essential at these locations to confer mechanical resistance and withstand strains encountered by cells (Acehan *et al.*, 2008; Weber *et al.*, 2012; Conway *et al.*, 2013).

*In vitro* assembly assays have demonstrated autonomous formation of intermediate filament networks that are dense, notoriously insoluble, and subject to little change (Sanghvi-Shah and Weber, 2017). Despite these observations, emerging evidence in the cellular environment has shown that these networks are, in fact, subject to ongoing modification (Vikstrom *et al.*, 1992; Windoffer *et al.*, 2004; Kolsch *et al.*, 2010). In conjunction with descriptions of ongoing filament network maintenance, IF networks have been shown to undergo dramatic and large scale reorganization in response to onset of mechanical stresses (Conway *et al.*, 2013). For instance, application of shear stress to cells results in both remodeling of cytoplasmic keratin IFs and an increase in dynamic exchange rate within these filaments (Sivaramakrishnan *et al.*, 2009). In our own previous work, we have shown that collectively migrating mesendoderm cells from the frog embryo demonstrate ongoing reorganization of keratin IFs to cell-cell contacts under tension (Weber *et al.*, 2012). Application of force to these cells through the cell-cell adhesion receptor C-cadherin induces a rapid shift in the cellular keratin network from widespread cytoplasmic to proximal to the cadherin junction experiencing tension. This distinct keratin recruitment presumably fortifies the mechanically sensitive junction and provides an instructive cue for directional cellular migration away from the site bearing mechanical load.

While several studies address the manner in which IFs are continually being formed (Windoffer *et al.*, 2004; Kolsch *et al.*, 2010), few to date have interrogated the processes that govern the disassembly and reorganization of IF networks. The phosphorylation status of IF proteins is a critical factor that determines features such as filament protein solubility, assembly and disassembly behaviors, and even aggregation of filament proteins (Ridge *et al.*, 2005; Snider and Omary, 2014). These modifications have cellular implications that include changes in signaling, cellular growth, and disease manifestation, among others. Several kinases and phosphatases have been shown to act on IF proteins (Ridge *et al.*, 2005; Omary *et al.*, 2006; Sivaramakrishnan *et al.*, 2009; Ju *et al.*, 2015). However, how phosphorylation and dephosphorylation alters intermediate filament conformation and protein binding partners is largely unknown. Moreover, a substantial gap also exists in identification of intermediate filament binding proteins that direct IF disassembly and/or recruitment of IFs to distinct cellular compartments.

The 14-3-3 proteins are a family of seven isoforms that have been demonstrated to bind to sequence motifs containing phosphorylated serine or threonine residues (Muslin *et al.*, 1996; Yaffe *et al.*, 1997). 14-3-3 homo and heterodimers have been shown to exert control over target substrates by alternatively limiting or enhancing affinity of these molecules for other binding partners (Tzivion *et al.*, 2000; Margolis *et al.*, 2006; Zhou *et al.*, 2010; Obsil and Obsilova, 2011). In this fashion, 14-3-3 proteins are recognized to modulate the activity of a variety of IFs (Li *et al.*, 2006; Miao *et al.*, 2013), as well as other cytoskeleton associated proteins such as motors and scaffold molecules (Jin *et al.*, 2004; Roberts *et al.*, 2013; Sehgal *et al.*, 2014; Vishal *et al.*, 2018).

We have previously demonstrated a mechanosensitive reorganization of the keratin network to cadherin mediated cell-cell contacts (Weber *et al.*, 2012). It is abundantly clear that the keratin network dynamics must be regulated to determine where and when such rearrangement events occur. We propose that 14-3-3 regulates the force triggered reorganization of the widespread keratin network to the cell-cell junction. In this study, we investigate a fundamental role for 14-3-3 interaction with Keratin 19 in modulating intermediate filament dynamic exchange and serving as a recruitment scaffold at cell-cell contacts. This interplay between the keratin network and 14-3-3 proteins may underlie signaling events that occur during collective cell migration, which in turn regulate the migratory behavior itself.

## Results

### 14-3-3 Protein Expression is Ubiquitous Throughout Early Embryonic Stages and Tissues

Previous work by Lau, Wu and Muslin examined mRNA expression levels and patterns of six different 14-3-3 isoforms during stages 2-38 of *Xenopus* embryonic development (Lau *et al.*, 2006). While mRNA expression was found to be abundant for 14-3-3 β, ɛ, and τ during stages 2-14, this is not necessarily indicative of protein expression in the embryo, which could be translationally regulated and/or maternally derived. To investigate 14-3-3 in the early *Xenopus* embryo, we thought it necessary to first examine 14-3-3 protein expression in developmental stages leading up to late gastrulation. Lysates were collected at time points beginning at one-cell stage through late gastrula (NF Stage 12.5), and immunoblot was performed using an antibody reactive with all isoforms of 14-3-3 (Figure 1A). Expression of 14-3-3 was detected throughout these early embryonic stages. Although 14-3-3 protein levels did increase slightly following mid-blastula transition (stage 7-9) when zygotic mRNA is transcriptionally upregulated, we saw relatively high and persistent levels of 14-3-3 proteins present throughout early development. To further examine the extent to which 14-3-3 was present in various embryonic tissues, gastrulating embryos were dissected into several regions (Figure 1B), and protein lysates were prepared. We examined the expression of 14-3-3 proteins in the animal cap, marginal zone, vegetal hemisphere, and mesendoderm tissues compared to that of whole embryos (Figure 1C). 14-3-3 was found to be present in all tissues of *Xenopus* gastrula. Although 14-3-3 protein levels varied between tissues, 14-3-3 expression was neither exclusive to nor absent from any one particular region, suggesting broadly ubiquitous functions for 14-3-3 across different tissue types. Relative to the housekeeping protein GAPDH, levels of 14-3-3 were particularly high in the collectively migrating mesendoderm tissue (Figure 1C).

**Figure 1.**
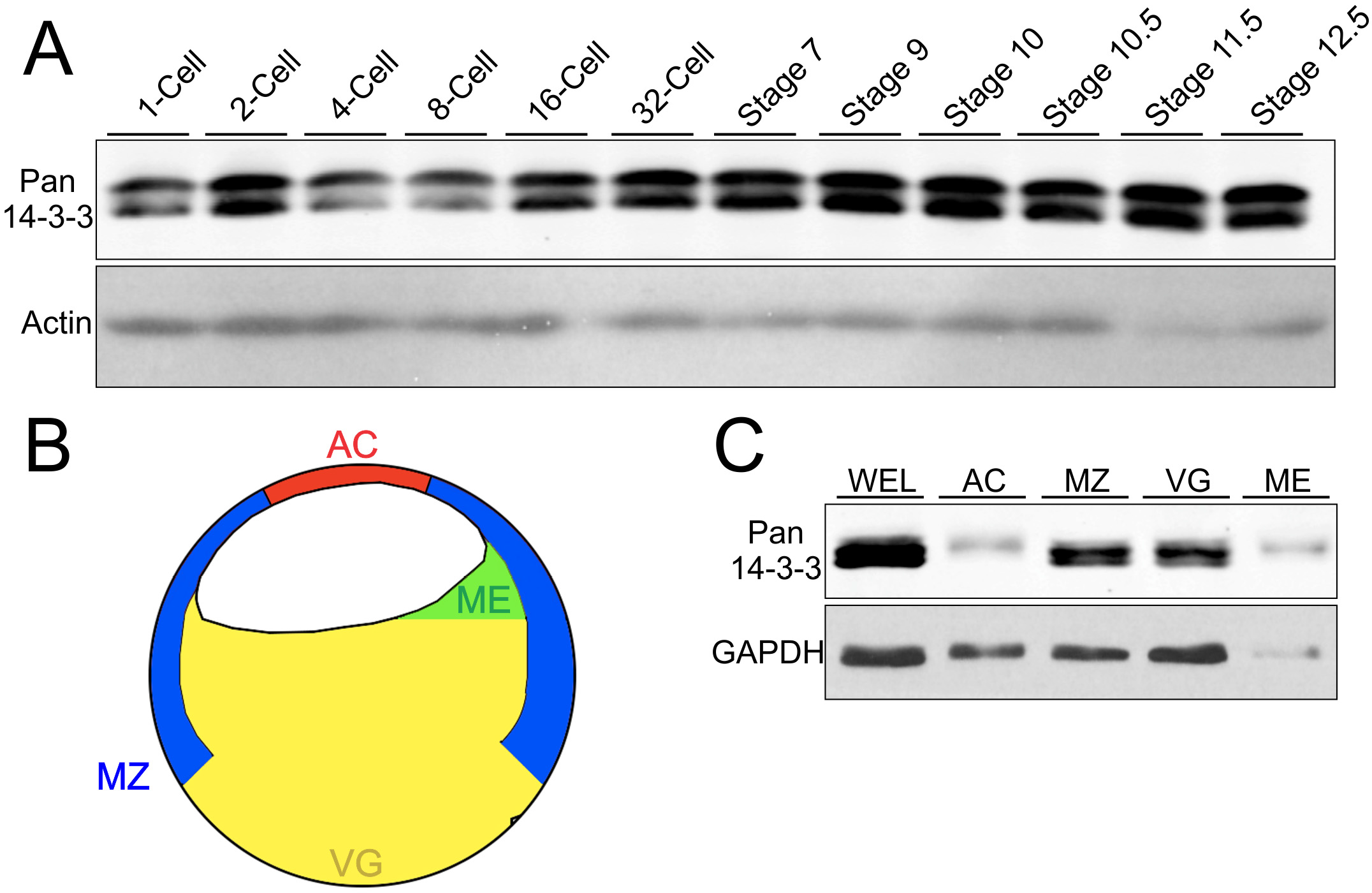
14-3-3 protein expression is ubiquitous across early embryonic stages and tissues. A) Whole embryo lysates were immunoblotted for 14-3-3 using a pan antibody that detects multiple isoforms. Each lane represents approximately 50 μg. B) Colored schematic of a bisected *Xenopus* embryo at gastrula depicting major tissue divisions. The tissues include the animal cap (AC), mesendoderm (ME), marginal zone (MZ), vegetal hemisphere (VG), and whole embryo lysate (WEL). C) Embryos were dissected into separate tissues and immunoblotted using pan 14-3-3 antibody to examine expression across the gastrulating embryo. Each lane represents approximately 50 μg.

### K19 Associates with 14-3-3 in Whole Embryos and Collectively Migrating Tissues

We next sought to identify specifically which 14-3-3 protein isoforms were present in gastrula and examine whether an association with keratin intermediate filament proteins could be detected. Co-immunoprecipitation was performed using pan-14-3-3 antibody with lysates from whole embryos or from mesendoderm tissue only. Samples were then separated by gel electrophoresis and stained to visualize protein band pattern (Figure 2A). Several distinct protein bands were visible in 14-3-3 immunoprecipitated lysates at the expected molecular weight for 14-3-3 proteins, between 25-30 kD. Additionally, an especially prominent band was detected at approximately 48 kD. These bands were selected for proteomic screening using LC/MS-MS.

**Figure 2.**
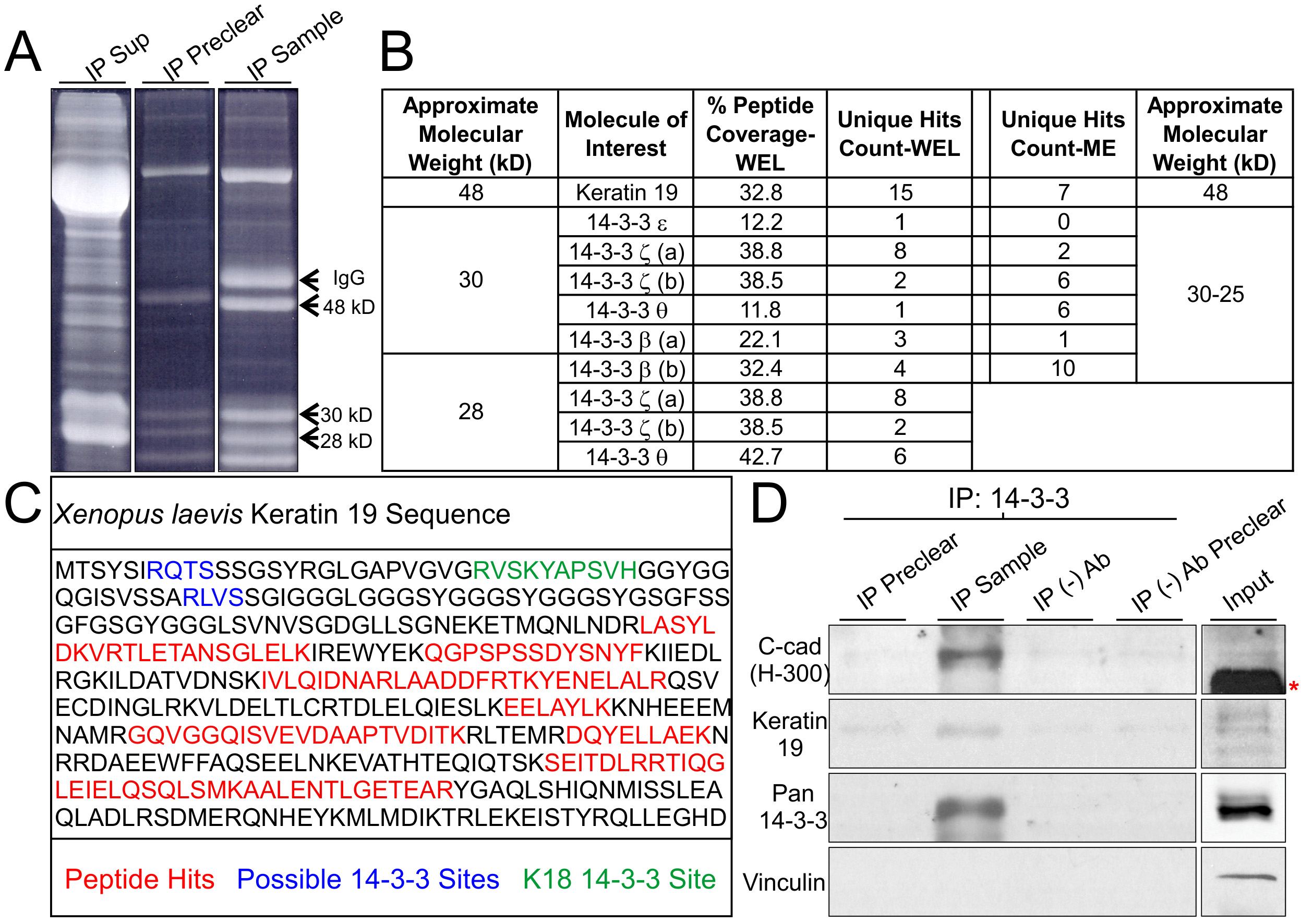
*Xenopus* Keratin 19 associates with 14-3-3 proteins and C-cadherin. A) Pan 14-3-3 immunoprecipitates from whole embryo lysates prior to band extraction and processing using LC/MS-MS. Prominent bands at 48, 30, and 28 kD were processed. Heavy chain IgG from the antibody used for IP was not excised. B) Table summary of relevant proteins detected in gel extracts processed using LC/MS-MS. Experiments were conducted using 14-3-3 immunoprecipitates from whole embryo lysates (WEL) as well as lysates from mesendoderm (ME) tissue only. Analysis was performed using Scaffold 4.7.3 C) Summary schematic of Keratin 19 peptides (red) detected in the 48 kD sample. Peptides are depicted within the context of the Keratin 19 primary structure and alongside described (green) and predicted (blue) possible 14-3-3 interaction sites. D) 14-3-3 proteins were immunoprecipitated from whole embryo lysates and immunoblotted for C-cadherin, Keratin 19 and Vinculin. The prominent band labeled by the asterisk is yolk protein.

Analysis of peptides identified by mass spectrometry confirmed the presence of multiple 14-3-3 protein isoforms (Figure 2B). Expanded datasets from the LC/MS-MS analyses can be found in Supplemental Figure 1. Several unique peptides as well as substantial total protein sequence coverage were found for 14-3-3 isoforms ζ, β, and τ in both whole embryo lysates and mesendoderm tissue alone (Figure 2B). Isoform 14-3-3ɛ was detected with lower unique peptide and coverage spectra than other isoforms and was not detected in mesendoderm. Interestingly in the analysis of the 48 kD band, Keratin 19 (K19) was detected with many unique peptides and substantial coverage in 14-3-3 co-immunoprecipitates from both *Xenopus* whole embryo lysates and mesendoderm (Figure 2C). It is important to note that K19 is not a keratin found in the epidermis of *Xenopus* nor humans, ruling out the possibility of this result occurring due to keratin contamination. To our knowledge, this is the first time K19 has been shown to associate with 14-3-3.

14-3-3 is known to bind protein substrates containing particular primary amino acid sequences. Notably the motif sequence RXXS/TP is a preferred target, but other variations do exist (Muslin *et al.*, 1996; Yaffe *et al.*, 1997; Johnson *et al.*, 2010). Several keratins, including K18, have been identified as phosphoserine-modified to become 14-3-3 substrates (Ku *et al.*, 1998). Using three different bioinformatics analyses, we examined whether K19 might also contain similar binding phosphorylated sites and motifs that could provide 14-3-3 docking sites (Supplemental Figure 2). K19 contains three major phosphorylated sites, S10, S33 and S52, which are highly conserved across phyla (Figure 2C and Supplemental Figure 2). S33 is of particular interest because it has previously been shown to be the primary phosphorylation site of K19 in several signaling and IF remodeling events (Zhou *et al.*, 1999; Ju *et al.*, 2015). We confirmed the association between 14-3-3 and K19 by immunoprecipitating 14-3-3 from whole embryo lysates and immunoblotting for K19 (Figure 2D). Indeed, K19 was present in 14-3-3 immunoprecipitated samples by immunoblot analysis as well.

The keratin intermediate filament network has been shown to be reorganized proximal to cell-cell adhesions in mesendoderm cells (Weber *et al.*, 2012). This reorganization of the keratin network occurs as a consequence of tugging forces on cell-cell interactions (Weber *et al.*, 2012). In *Xenopus* gastrula, the classical cadherin protein C-cadherin provides the primary means of cell-cell adhesion (Heasman *et al.*, 1994). Since keratin intermediate filaments are recruited to C-cadherin in mesendoderm cells as a function of applied force on the adhesions, we next investigated whether 14-3-3 was also associated with C-cadherin. 14-3-3 co-immunoprecipitated lysates were positive when immunoblotted for C-cadherin (Figure 2D). Interestingly, although vinculin has been implicated in mechanosensation in both cadherin cell-cell adhesions and focal adhesions by several reports (Riveline *et al.*, 2001; le Duc *et al.*, 2010), we did not find vinculin to be associated with 14-3-3 (Figure 2D).

### Keratin Filaments Co-localize with 14-3-3 at Cell Boundaries

Because 14-3-3 was found to biochemically associate with K19, we performed immunofluorescence to assess the relationship between 14-3-3 and keratin filament networks *in situ*. To determine the localization of 14-3-3 proteins in mesendoderm, immunofluorescence was performed on sagittal sections of gastrulating *Xenopus* embryos. We found that several tissues showed compartmentalization of 14-3-3 rather than diffuse cytoplasmic distribution. In mesendoderm, 14-3-3 proteins localized clearly and distinctly to cell boundaries at lower magnifications (Supplemental Figure 3). At higher magnifications, we were able to more precisely assess the subcellular distribution of 14-3-3, especially in relation to keratin filaments.

Keratin filaments demonstrate force-dependent localization to areas of cell-cell contact in collectively migrating mesendoderm (Weber et al. 2012). As expected, keratin filaments were found localized to the posterior of migratory mesendoderm cells, where they link to cell-cell contacts (Figure 3A). 14-3-3 shows strong co-localization with these densities of keratins present at points of cell-cell adhesion (Figure 3B,C). Similar co-localization was also seen in coimmunofluorescence labeling of mesendoderm explants (Figure 3E-G). Regions of signal overlap are filamentous in shape in both sagittal whole embryo sections and explanted tissue (Figure 3C, E’-G’). Rather than extending across the entire filament density, the co-labeling occurs specifically at the area proximal to the cell-cell adhesion. 14-3-3 also appeared to exhibit concentrated intensity in lamellipodia of mesendoderm cells, where keratin filaments are notably absent. Interestingly, 14-3-3 labeling in lamellipodia lacked the filamentous pattern observed in the posterior of cells.

**Figure 3.**
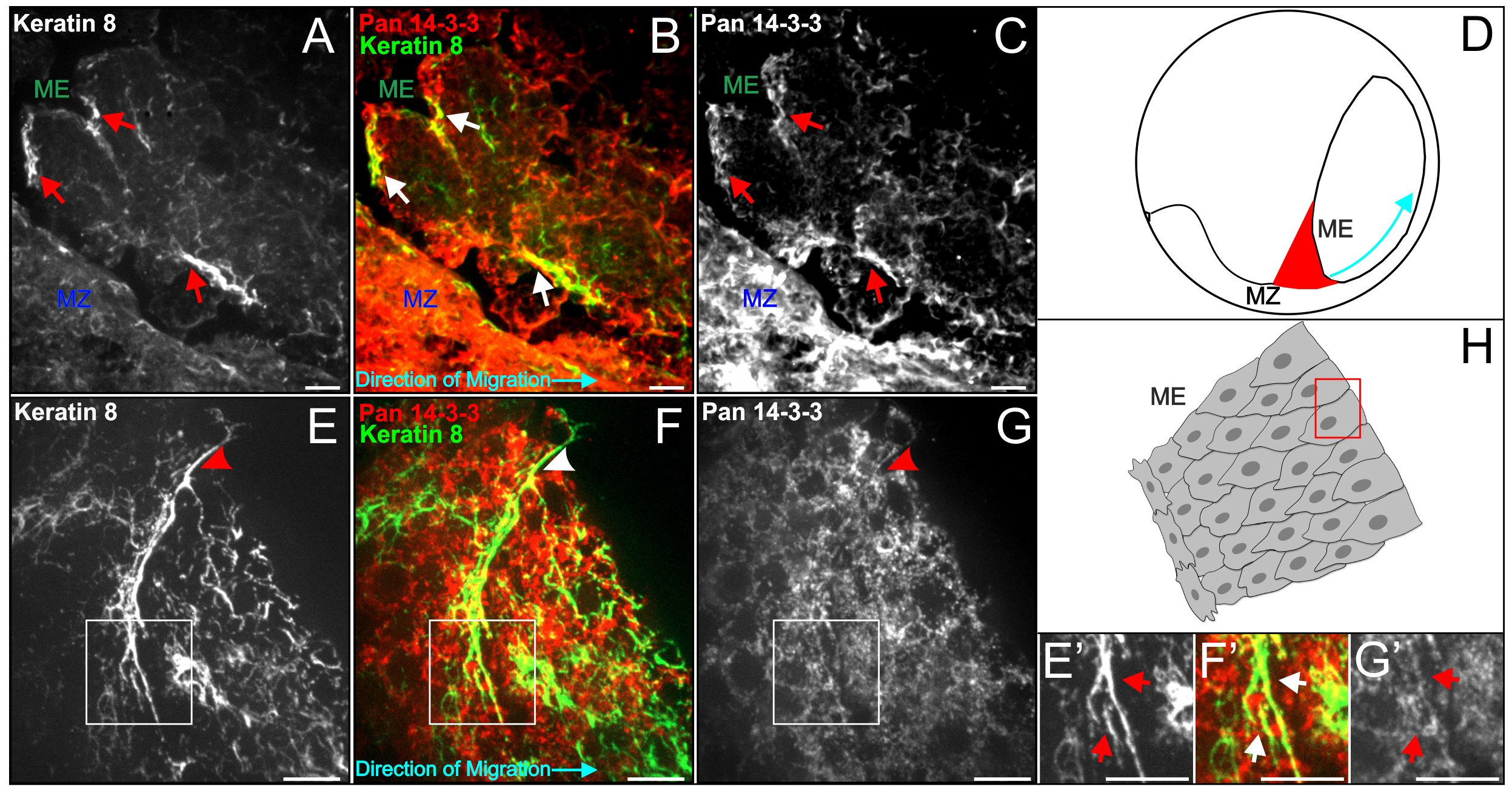
Filaments recruited to cell-cell adhesions associate with 14-3-3. A-C) Sagittal perspective of a cryosectioned gastrulating embryo labeled immunocytochemically for 14-3-3 proteins (red) and Keratin 8 (green). Areas where filamentous co-localization was detected in panel B are illustrated by arrows in all three panels. D) Embryo schematic depicting orientation of cells in A-C relative to the sagittally sectioned embryo. Blue arrow indicates direction of tissue migration. E-H) Leading edge of a mesendoderm explant demonstrating association between 14-3-3 and Keratin 8 at a cell-cell interface (arrowheads). Closer inspection of the keratin morphology at this area (E’-G’) reveals filamentous 14-3-3 labeling. H) Explant schematic depicting the cell pair (E-G) relative to the rest of the tissue. Images are z-stacks (maximum intensity projection). Scale bars are 10 μm.

### 14-3-3 Proteins are Distributed Proximal to Cell-Cell Adhesions

C-cadherin co-localizes with keratin filaments at points of cell-cell contact in mesendoderm and co-immunoprecipitates with keratin in *Xenopus* gastrula (Weber *et al.*, 2012). Given that co-immunoprecipitation of 14-3-3 also shows association with C-cadherin, we examined how the subcellular localization of each relate to one another. Curiously, although 143-3 appears at the cell periphery at lower magnification (Supplemental Figure 3), an appreciable separation between 14-3-3 and C-cadherin is evidenced at higher magnification (Figure 4A-D). 14-3-3 is increasingly concentrated nearer to cell-cell contacts (Figure 4E-G) as compared to more centrally located regions of the cytoplasm. However, at the more precise point of cell-cell contact, indicated by C-cadherin-eGFP labeling, we noted a decrease in 14-3-3 signal intensity (Figure 4H). Taken together with evidence of biochemical association, this subcellular space suggests that the association between 14-3-3 and C-cadherin in the keratin-cadherin complex is indirect. 14-3-3 appears to localize with keratin intermediate filaments and concentrate near cell-cell contacts, but not exactly at the cell-cell contact itself.

**Figure 4.**
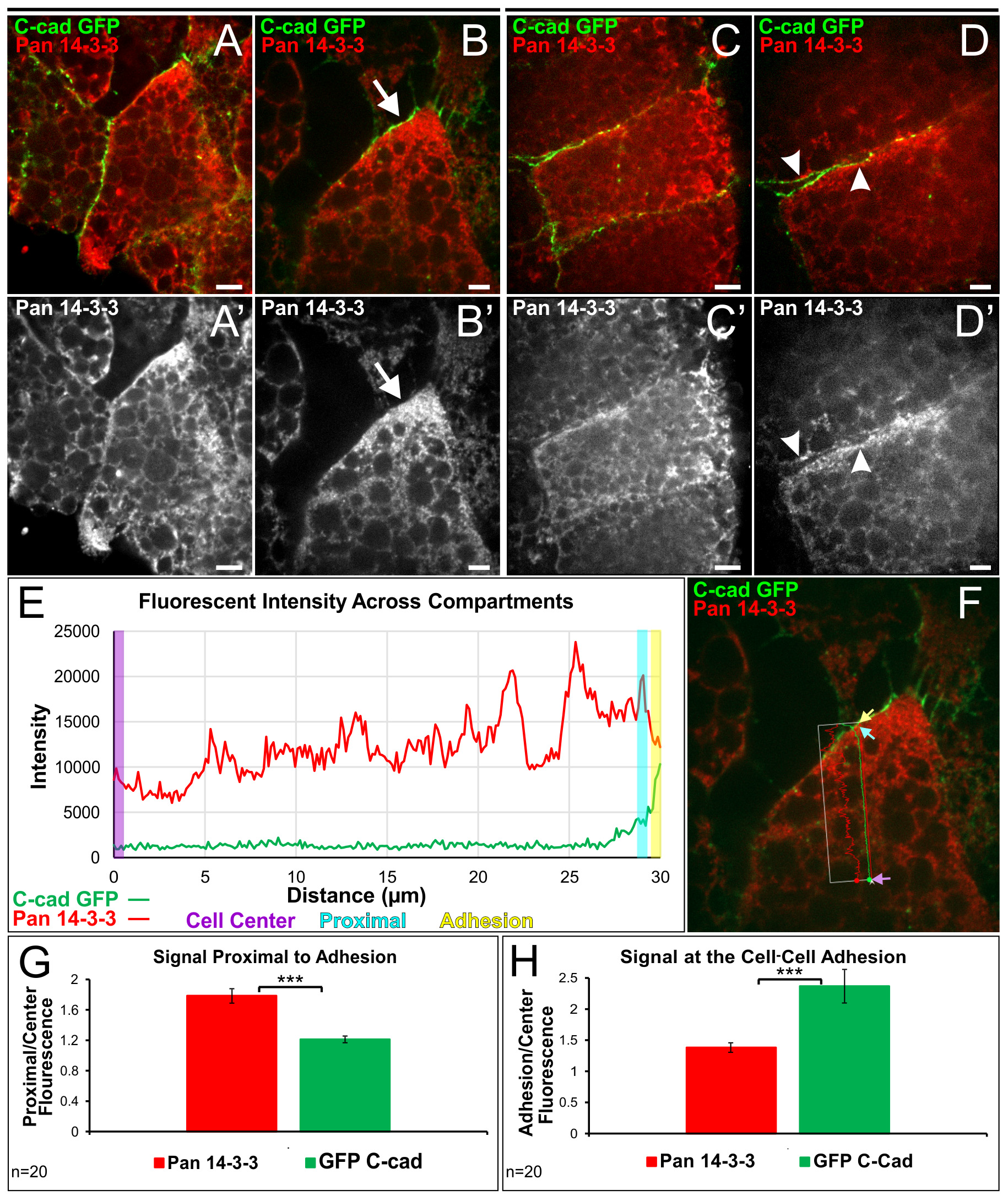
14-3-3 proteins are distributed proximally to cell-cell adhesions. A-B’) Immunofluorescent image of a leading edge mesendoderm cell labeled for 14-3-3 and expressing C-cadherin (eGFP label). Increased magnification was utilized to capture the rear contact of the cell (arrow, B and B’). The direction of migration in A-B’ is down. C-D’) Image of a leading edge mesendoderm cell highlighting a rear-lateral contact (arrowheads), with D and D’ at higher magnification. The direction of migration in C-D’ is left. E) Representative linescan analysis of fluorescence across cellular compartments in panel B. Lines extended from the approximate middle of the cell to the area just prior to C-cadherin signal and to the onset of C-cadherin signal. Measurements were taken from the cell center (purple rectangle), proximal to the adhesion (teal rectangle), and at the adhesion (yellow rectangle). Each rectangle represents 0.5 μm in length. F) Linescan measurement of panel B. Colored arrows indicate region represented by rectangles in E. G and H) Comparison of the mean increase in 14-3-3 signal and C-cadherin signal from the cell center to the area proximal to the cell-cell adhesion (in panel G) and at the cell-cell adhesion (in panel H). Analysis was performed using paired sample t-tests, *** = p<0.001. Error Bars are ± SEM. Scale bars are 10 μm (A,A’,C,C’) and 5 μm (B,B’,D,D’).

### Intermediate Filament Dynamic Exchange is Disrupted by 14-3-3 Inhibition

The observation that 14-3-3 associates with keratins and co-localizes with filaments implies a shared functional relationship. Given that 14-3-3 proteins have been shown to have a role in disassembly of IFs (Li *et al.*, 2006; Miao *et al.*, 2013) and to associate with keratin filaments induced to remodel by okadaic acid treatment (Strnad *et al.*, 2002), one possible function of this association is to facilitate modification of pre-existing filament networks. Modification of polymerized filaments involves addition or subtraction of IF proteins along the length of a filament polymer, known as lateral exchange or dynamic exchange (Vikstrom *et al.*, 1992; Colakoğlu and Brown, 2009; Nöding *et al.*, 2014). Keratin networks show filament remodeling and 14-3-3 dependent increases in dynamic exchange rate after exposure to shear stress (Sivaramakrishnan *et al.*, 2009). Since changes in mechanical stress are continually transduced across junctions during collective cell migration and result in keratin reorganization (Weber *et al.*, 2012), we examined whether 14-3-3 might have a role in the keratin filament organization in mesendoderm explants. To functionally inhibit 14-3-3, we expressed a short peptide sequence (R18) previously shown to bind 14-3-3 with high affinity and block its binding to endogenous substrates (Petosa *et al.*, 1998; Wang *et al.*, 1999). A negative control peptide (R18M) containing two point mutations was used for comparison. We created mCherry-tagged R18 and R18M constructs so that we could readily identify living cells in which R18 or R18M was expressed. The ability of mCherry-R18, but not mCherry-R18M, to bind 14-3-3 was confirmed by co-immunoprecipitation analyses (Figure 5E).

**Figure 5.**
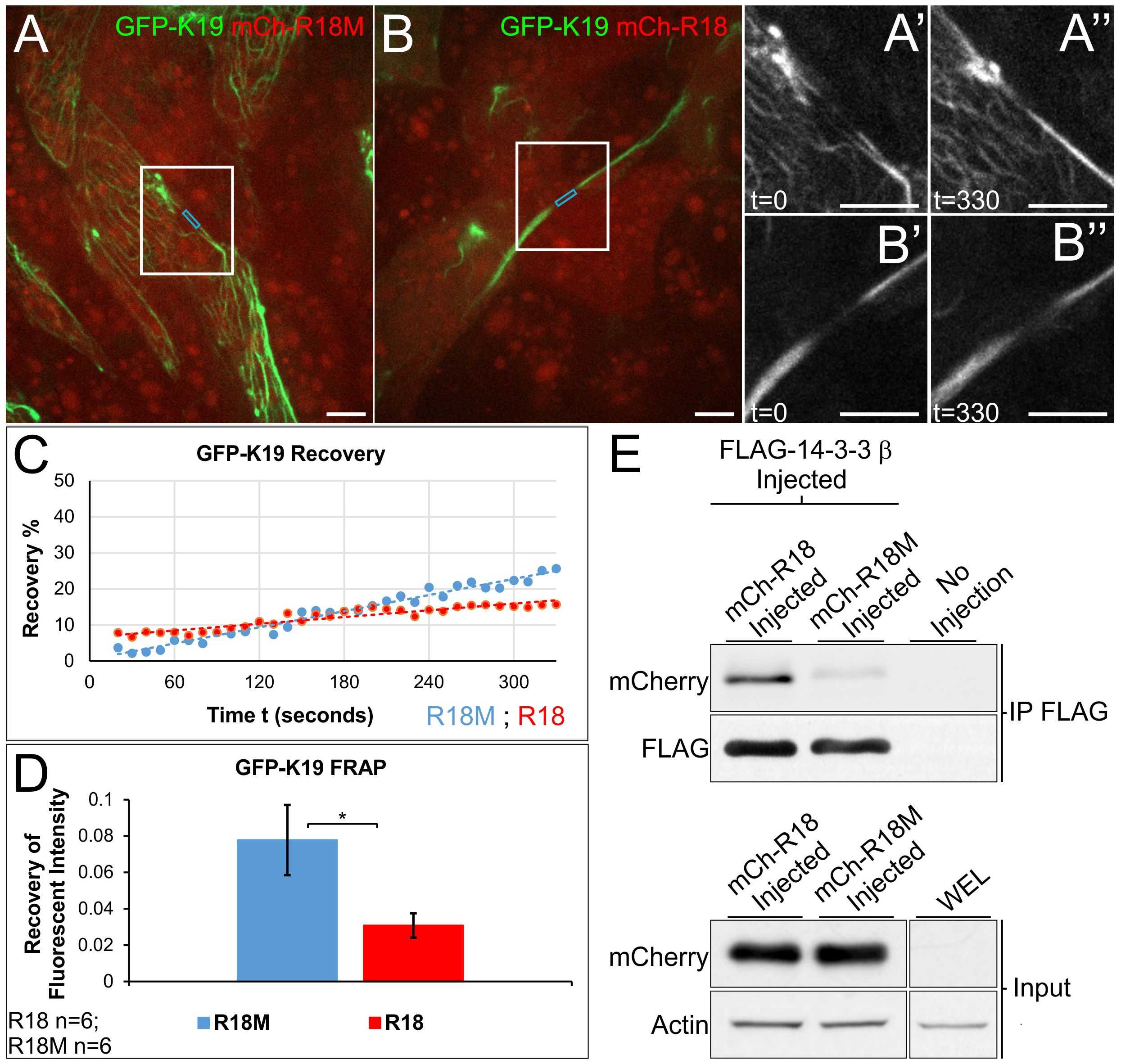
Mesendoderm Keratin filament dynamic exchange is decreased by 14-3-3 inhibition. A,B) Still images from photobleach and recovery time lapse movies (Supplemental Movie 1). Mesendoderm explants expressing either 14-3-3 inhibitor peptide mCherry-R18 or control peptide mCherry-R18M (red) with eGFP-K19 (green) were exposed to GFP photobleaching and fluorescent recovery at the site was measured. A’-B’’) Enlarged view of the region of filament bleaching (white boxes) during recovery measurements. The time annotations in seconds refer to start of capture (t=0) and end of capture (t=330). C) Representative analysis plotting fluorescent recovery against time of image capture. D) Comparison of mean eGFP-K19 recovery rate in explants expressing either mCherry-R18 or mCherry-R18M. Analysis was performed using a one tailed t-test, * = p<0.05. Error Bars are ± SEM. E) Co-immunoprecipitation performed using stage 10.5 lysates expressing FLAG-14-3-3P with either mCherry-R18 or mCherry-R18M. Scale bars are 10 μm.

To examine the role of 14-3-3, we co-expressed mCherry-R18 or R18M with eGFP-K19. R18 expression did not initially result in any apparent changes to the keratin network detectable by fluorescent microscopy. Through early gastrulation and explant preparation, no gross morphological changes were observed in the overall keratin network or subcellular localization of filaments. We suspect that this may be because of a delayed onset of R18 expression until after mid-blastula transition. Nonetheless, subtler effects on the intermediate filament dynamics were observed in native mesendoderm explants. FRAP experiments of eGFP-K19 revealed that 14-3-3 inhibition results in attenuation of dynamic exchange rate when compared to that of explants expressing control peptide R18M (Figure 5A-C). Despite inherent differences in the width and subcellular location of filaments bleached across trials, the lower recovery rate in 143-3 inhibited samples persisted (Figure 5C-D). Though additional reports provide evidence of different times of lateral exchange onset that range from within an hour to eight hours (Colakoğlu and Brown, 2009; Nöding *et al.*, 2014), we detect more rapid differences in recovery after 14-3-3 inhibition that occurs within minutes (Figure 5). This finding demonstrates that the interaction between 14-3-3 and Keratin 19 is required for modification of filaments that occurs on an appreciably fine timescale in motile tissues.

### 14-3-3 is Required for Keratin Recruitment to Cell-Cell Adhesions

Force transmitted through mesendoderm cell-cell contacts induces both changes in the keratin network throughout the cell and recruits filaments to sites of transduced tension (Weber *et al.*, 2012). We were surprised to find that expression of R18 did not dramatically alter the distribution of keratin intermediate filaments in the mesendoderm given its association with keratin and role in dynamic exchange. We reasoned that this could be because expression of R18 did not occur until after relatively stable cell-cell adhesions were formed. To determine whether 14-3-3 is required for filament recruitment at cell junctions, mesendoderm tissue expressing mCherry-R18 or R18M and eGFP-K19 was dissociated into single cells, plated on fibronectin substrate, and imaged as cells collided and formed cell pairs. Our work and others’ have shown that many migratory cells polarize in opposite directions upon collision, resulting in substantial force at the cell-cell junction (Nelson and Chen, 2003; Maruthamuthu *et al.*, 2011; Weber *et al.*, 2012; Mertz *et al.*, 2013). We additionally showed that keratin intermediate filaments are recruited to cell-cell adhesions between mesendoderm cells as a function of force on C-cadherin (Weber *et al.*, 2012). This was also the case in mesendoderm cells expressing the control construct mCherry-R18M (Figure 6A-C). When 14-3-3 was inhibited by expression of mCherry-R18, a distinct zone lacking fluorescent filaments was observed at points of *de novo* cell-cell adhesion (Figure 6D-F). This keratin-deplete zone was observed at a significantly higher frequency in 14-3-3 inhibited cells after establishment of cell-cell contact relative to controls (Figure 6). These regions lacking keratin IFs had a size range of 2 μm to 17 μm with an average of 7 μm when measured from the cell-cell contact to the onset of visible filament fluorescence. The zone was surprisingly uniform along the width of the cell-cell contact in these cells and very few R18-positive cells were found to have keratin recruited to cell-cell contacts (Figure 6G).

**Figure 6.**
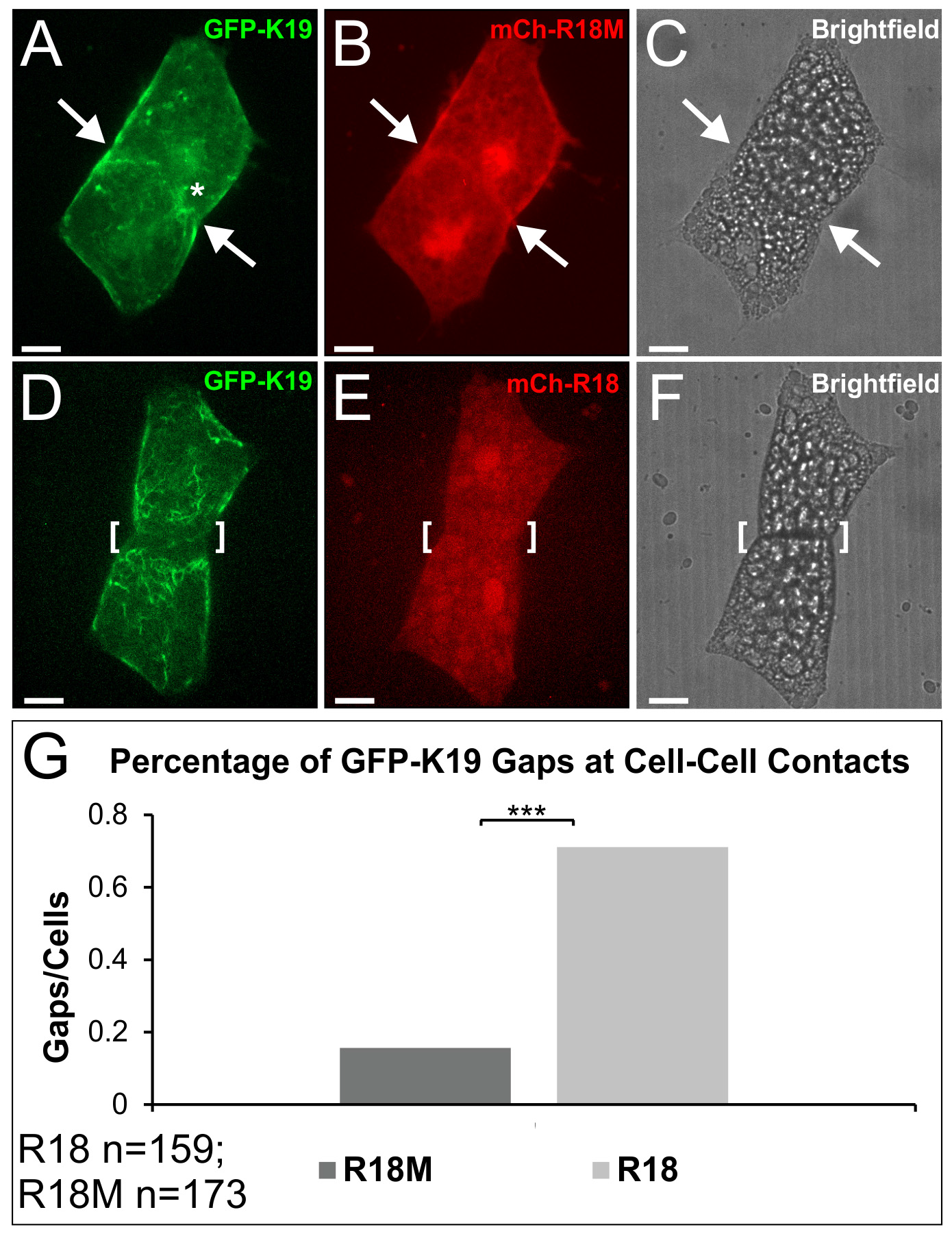
14-3-3 is necessary for recruitment of keratin to cell-cell contacts. A-C) Mesendoderm cell pair establishing *de novo* cell-cell contact after collision. Cells are expressing mCherry-R18M and eGFP-K19. The arrowhead depicts the cell-cell adhesion where keratin densities (asterisk) have localized. Image stack is 35 slices (10.54 μm). D-F) Post collision mesendoderm cell pair expressing mCherry-R18 and eGFP-K19. The bracket depicts the cell-cell adhesion that demonstrates a gap where keratin filaments have failed to reorganize. Image stack is 40 slices (10.92 μm). G) Comparison of the number of keratin gaps at cell-cell adhesions in post collision pairs expressing either mCherry-R18M or mCherry-R18. Analysis was performed using a z-test for proportions from two samples, *** = p<0.001. Fluorescent images are z-stacks (maximum intensity projection). Brightfield images are single planes. Scale bars are 20 μm.

### 14-3-3 Association with K19 is Sufficient for Cell-Cell Adhesion Targeting

Since inhibition of 14-3-3 prevented keratin recruitment to cell-cell adhesions, we wanted to see whether forced association of keratins with 14-3-3 would be sufficient to drive their targeting to cell junctions. 14-3-3 endogenously binds to phosphorylated serine and threonine residues contained in motifs within certain substrates (Yaffe *et al.*, 1997; Johnson *et al.*, 2010). Mutation of serine residues to non-phosphorylatable alanines (e.g. S33A) eliminates binding between keratins and 14-3-3; however, mutation of serine to phosphomimetic amino acids (e.g. S33D) is not sufficient to promote association between 14-3-3 and keratins, presumably because of differences in the side chain compared to phosphate groups (Ku *et al.*, 1998). In order to promote the association of K19 with 14-3-3, we created an eGFP-R18-K19 fusion protein (Figure 7A). Since R18 is a high affinity peptide for 14-3-3, this fusion protein would result in a traceable K19 protein with a constitutive signaling site for 14-3-3 binding that is phosphorylation independent.

**Figure 7.**
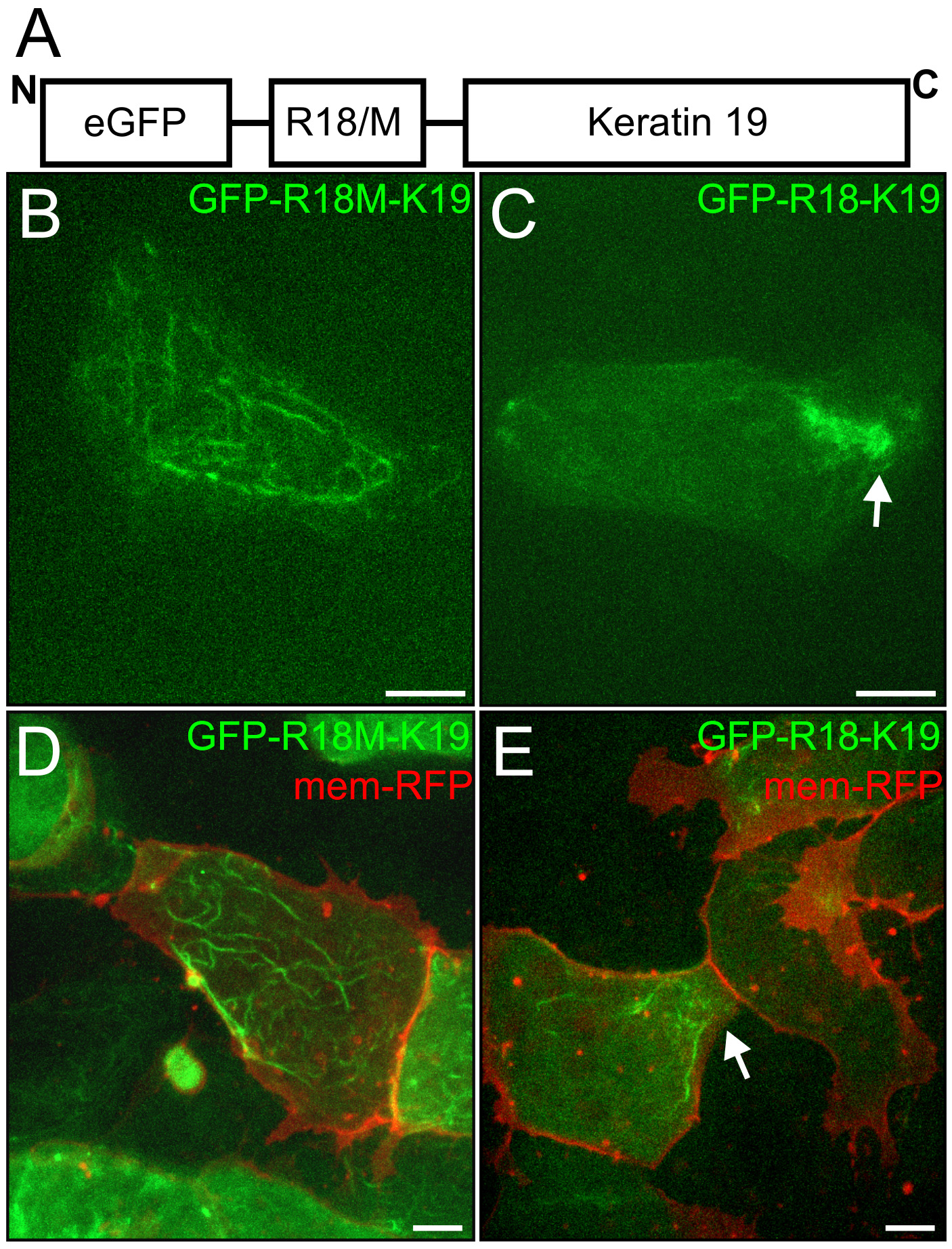
14-3-3 proteins induce recruitment of keratins to cell-cell adhesions. A) Schematic of fusion peptides created by the addition of R18 or R18M (R18/M) into eGFP-K19. B and C) Explanted mesendoderm cells mosaically expressing eGFP-R18M-K19 (in panel B) or eGFP-R18-K19 (in panel C). D and E) Explanted mesendoderm co-expressing mem-RFP and eGFP-R18M-K19 (in panel D) or eGFP-R18-K19 (in panel E). Arrows indicate areas where filament densities have localized. Images are z-stacks (maximum intensity projection). Scale bars are 10 μm.

Mosaic expression of these fusion constructs was established in mesendoderm tissue by injecting DNA, and the intermediate filament distribution in live mesendoderm cells, ~4 rows back from the leading edge, was examined by confocal microscopy. Expressed eGFP-R18M-K19 in mesendoderm explants forms filaments spread throughout the cell body but absent from cell protrusions (Figure 7B,D), typical of keratin networks described previously in unpolarized mesendoderm (Weber *et al.*, 2012). In mesendoderm expressing eGFP-R18-K19, the fluorescence appeared as densities that preferentially localize to areas of cell-cell adhesion and lack widespread expression throughout the cytosol (Figure 7C,E). In the most robust instances, eGFP-R18-K19 localized as an intense aggregate exclusively at the cell-cell boundary (Figure 7C). In other cases, distinct filaments could be seen and the population was limited to near the cell-cell adhesion (Figure 7E). The forced association of K19 with 14-3-3 through this fusion construct created a distribution that in many ways was the opposite of when 14-3-3 was inhibited (compare Figure 7C with Figure 6D). Altogether these data indicate a role for 14-3-3 association with K19 in the preferential targeting of keratin to cell-cell adhesions.

## Discussion

Our findings in the current study lead us to propose that 14-3-3 proteins are responsible for targeting keratin intermediate filaments to sites of cell-cell adhesion that are transmitting tension. Expression of a fluorescent keratin fused to a high affinity 14-3-3 binding domain causes filaments assembled from these keratins to mobilize to cell-cell junctions. In reciprocal support, inhibition of 14-3-3-through expression of a peptide inhibitor results in the failed recruitment of keratin filament to sites of *de novo* cell-cell adhesion. These observations provide evidence for the first time that 14-3-3 functions as a critical spatial determinant of the intermediate filament network in response to mechanical loading at cell contacts.

Cytoskeletal structures must be able to efficiently sense tension and undergo remodeling in order to fortify cells against changes in the direction, magnitude, and nature of tension transmitted. Rapid changes in force application require concomitant responses by cytoskeletal proteins, necessitating mechanistic filament assembly strategies that are effective within a short time scale. Cellular mechanisms that induce changes within a shorter time-scale than that required for biosynthesis include pathways that make post-translational modifications to preexisting proteins. Intermediate filament proteins, which have no known intrinsic catalytic activity, integrated within a filament network are modified by other molecules that respond to changes in cellular tension, such as kinases (Loschke *et al.*, 2015). It follows that accessory proteins recognize and bind these post-translational modifications and accordingly alter the filament dynamics and spatial localization of intermediate filament proteins. These interactions can recruit filaments to particular locales within the cell and change polymerization-depolymerization characteristics. Our data support a functional role for 14-3-3 in mediating the recruitment to cell-cell adhesions and dynamic exchange of keratin intermediate filaments in cells that are actively migrating and encountering changes in the direction and magnitude of loading forces.

We show here an interaction between 14-3-3 and keratin proteins. Indeed, 14-3-3 has been demonstrated to interact with a variety of intermediate filaments (Tzivion *et al.*, 2000; Li *et al.*, 2006; Miao *et al.*, 2013), including some keratin types (Ku *et al.*, 1998; Kim *et al.*, 2006; Margolis *et al.*, 2006; Sivaramakrishnan *et al.*, 2009; Boudreau *et al.*, 2013). We show for the first time an interaction between K19 and 14-3-3. Previous work has shown that K18 directly binds to 14-3-3 when phosphorylated on serine 33, but an association between K19 and 14-3-3 was not observed (Ku *et al.*, 1998). Nonetheless, this serine residue is conserved between K18 and K19 and is a major phosphorylation target in K19 (Ku *et al.*, 1998; Zhou *et al.*, 1999; Ju *et al.*, 2015). 14-3-3 associates with a number of Type I acidic keratins aside from K18 (Kim *et al.*, 2006; Boudreau *et al.*, 2013). Serine residues 10, 33 and 52 of K19 are predicted to be phosphorylated by several kinases including protein kinase C ζ and to be 14-3-3 binding sites by our bioinformatics analysis (Supplemental Figure 2). Regardless of which specific phosphorylation site provides for interaction with 14-3-3, an association is evident by our experimental observations and supported by numerous predictive computational analyses. While one might contend that K19 could be associating with 14-3-3 indirectly through other keratins, K19 was uniquely abundant in our 14-3-3 immunoprecipitation samples.

What is the interaction of 14-3-3 with K19 actually doing to the intermediate filaments? At a whole-cell level, association of keratins with 14-3-3 is necessary for keratin network disassembly during mitosis (Liao and Omary, 1996). Knockdown of 14-3-3 has been shown to result in abnormal increases in keratin filament abundance and decreases in soluble filament precursors (Boudreau *et al.*, 2013). Fluid flow shear stress of alveolar cells results in phosphorylation of known 14-3-3 sites in keratin 18 along with increases in dynamic exchange that are 14-3-3 dependent (Sivaramakrishnan *et al.*, 2009). Similarly, we see decreased FRAP recovery when 14-3-3 is inhibited in eGFP-K19 expressing mesendoderm tissue, which is known to have intrinsic mechanical stresses through the cell-cell contacts (Weber *et al.*, 2012; Sonavane *et al.*, 2017). Expression of R18 peptide results in inhibition of 14-3-3 in multiple contexts throughout the cell rather than perturbation of the subset of 14-3-3 molecules that associate with the keratin-cadherin complex. Nonetheless, we find it remarkable that 14-3-3 inhibition still resulted in a distinct change in keratin subcellular distribution as a consequence of cell-cell adhesion signals.

We find that 14-3-3 provides for targeting of K19 to cadherin-mediated cell-cell contacts. Forced association of K19 with 14-3-3 localized this population of keratin filaments to cell-cell contacts, and conversely, inhibition of 14-3-3 prevented the recruitment of keratin to these junctions. 14-3-3 is observed decorating the filamentous keratin population proximal to these adhesions. Despite finding biochemical association between 14-3-3 and C-cadherin, we note that at high resolution these molecules do not precisely co-localize. This suggests that there is a scaffold intermediate that may facilitate interaction between 14-3-3/keratin protein complexes and other molecules of the cell-cell junction, and which may play a role in the reorganization event. 14-3-3 has been demonstrated to associate with a number of molecules important for the linkage of cytokeratins to the cell-cell adhesion (Acehan *et al.*, 2008) including plakoglobin (Sehgal *et al.*, 2014; Vishal *et al.*, 2018) and plakophilin (Jin *et al.*, 2004; Roberts *et al.*, 2013). The association and function of 14-3-3 in relation to keratin intermediate filaments may provide an interface between cytoskeletal networks and a host of scaffolding proteins and signal transduction pathways (Margolis *et al.*, 2006; Loschke *et al.*, 2016; Sonavane *et al.*, 2017). In turn, the local recruitment of keratin at sites of tension due to local signaling/phosphorylation events could promote a positive feedback loop for keratin filament assembly dynamics (Ridge *et al.*, 2005; Woll *et al.*, 2007; Ju *et al.*, 2015; Sonavane *et al.*, 2017). The enriched presence of 14-3-3 near both the cell-cell adhesion and in the lamellipodia suggest either non-intermediate filament related functions in the latter region or a possible function in intermediate filament formation (Windoffer *et al.*, 2004, 2011). Further elucidation of these macromolecular complexes in polarized migratory cells will be the subject of ongoing studies.

The significance of K19 association with 14-3-3 to development of the early embryo remains uncertain. Previous work has shown important roles for 14-3-3 in *Xenopus* development through the targeted knockdown of different isoforms by morpholino oligonucleotides (Lau *et al.*, 2006). Knockdown of certain 14-3-3 isoforms caused severe gastrulation defects including exogastrulation and failed mesodermal patterning (Lau *et al.*, 2006). Knockdown or inhibition of K8, the lone Type II keratin in early embryogenesis, similarly induces exogastrulation (Klymkowsky *et al.*, 1992; Torpey *et al.*, 1992). The specific functions of K18 and K19 in early development have yet to be determined, but it is likely that there exists some functional redundancy as evidenced by murine knockouts (Magin *et al.*, 1998). It is notable that K19 does not have a tail domain, making it structurally deviant from other IF proteins (Herrmann *et al.*, 1996; Kirmse *et al.*, 2007; Lee *et al.*, 2012). This has been speculated to result in keratin networks with greater dynamic potential (Hofmann and Franke, 1997; Fradette *et al.*, 1998) and are thus suited to a dynamic tissue such as collectively migrating mesendoderm.

Molecular control of reorganization of filaments is a process that is as critical to cellular migration as it is to cellular and junctional integrity. The results of the current study lend insights into the mechanisms of active tissues ranging from cellular sheets in migration to three dimensional tissues that balance changes in loading (Winklbauer *et al.*, 1992; Weber *et al.*, 2012). These observations can also inform cancer models that detect roles for 14-3-3 proteins and keratins in promoting invasiveness of cells (Boudreau *et al.*, 2013; Cheung *et al.*, 2013; Deng *et al.*, 2013; Ju *et al.*, 2015). Decrease in the ability of IFs to form the networks that fortify sealed junctions across cells, enable cells to form resistant tissues, and coordinate the activity of tissues has implications for essential processes including barrier function, embryonic tissue patterning, would healing, and cancer progression‥

## Materials and Methods

### Xenopus Embryo In Vitro Fertilization and Staging

Embryos were obtained and cultured using standard methods and staged according to (Nieuwkoop and Faber, 1994). Embryos were dejellied in 2% cysteine and cultured at 15°C in 0.1X Modified Barth’s saline (MBS; 1X MBS: 88 mM NaCl, 1 mM KCl, 2.5 mM NaHCO_3_, 0.35 mM CaCl_2_, 0.5 mM MgSO_4_, 5 mM HEPES pH 7.8).

### Xenopus Embryo Protein Lysate Preparation and Western Blot

*Xenopus* whole embryos were solubilized in lysis buffer (50 mM Tris-HCl pH 7.5, 1 mM phenylmethylsulfonofluoride (PMSF), 1% mammalian protease inhibitor cocktail (Sigma-Aldrich, P2714), 0.2 mM H_2_O_2_, 3 mM sodium pyrophosphate, 1 mM sodium orthovanadate, 10 mM sodium fluoride, sodium β-glycerophosphate [10 mg/ml], 1% Triton X-100 (Sigma-Aldrich, T-9284) or 1% Tergitol type NP-40 (Spectrum Biosciences, T1279). Protein samples were prepared in 2x Laemmli buffer supplemented with 5% β-mercaptoethanol, incubated on a 95°C heat block for five minutes, and loaded onto 12% SDS-PAGE gels. Proteins were transferred onto nitrocellulose membrane prior to incubation with antibodies.

### Antibodies

Several antibodies were used throughout this study: pan 14-3-3 pAb (Santa Cruz, K19 sc-629), 1h5 (Keratin 8) mAb, pan Cadherin pAb (Santa Cruz, H-300 sc-10733), Vinculin mAb (Millipore, MAB3574), Keratin 19 mAb (Progen, 61010), Actin-HRP (Sigma-Aldrich,A3854), GAPDH mAb (Abcam, mAbcam 9484), GFP mAb (Invitrogen, A-11120), mCherry pAb (BioVision, 5993-100), and FLAG mAb (Sigma-Aldrich, F1804). The 1h5 anti-XCK1(8) monoclonal antibody developed by Michael Klymkowsky was obtained from the Developmental Studies Hybridoma Bank developed under the auspices of the NICHD and maintained by The University of Iowa, Department of Biology, Iowa City, IA 52242.

### Plasmids

The following plasmids were used in this study: pCS2-mCherry-R18, pCS2-mCherry-R18M, pCS2-eGFP-K19, pcDNA3.1-FLAG-HA-14-3-3-β (Addgene plasmid # 8999), pCS2-eGFP-C-cadherin (B. Gumbiner, University of Washington), pCS2-eGFP-R18-K19, pCS2-eGFP-R18M-K19, pCS2-mem-RFP. Oligonucleotides that translate to the described R18 and R18M primary sequences (Wang *et al*. 1999) were commercial synthesized (Genewiz) and cloned into pCS2 vector backbone by the author. *Xenopus laevis* K19 (NP_001084992.1) was obtained as full length cDNA (GE Dharmacon MXL1736-202774753) and cloned into pCS2 vector backbone by the author. 1478 pcDNA3 flag HA 14-3-3 beta was a gift from William Sellers (Addgene plasmid # 8999). The mem-RFP construct was a kind gift from Megason and Fraser (Megason and Fraser, 2003).

### RNA constructs and microinjection

RNA was prepared via *in vitro* transcription (Promega, P1420). DNA or RNA was diluted to microinject concentrations of 375-500 pg in 5 nl pulses for the following constructs: pCS2-eGFP-K19, pCS2-eGFP-R18-K19, and pCS2-eGFP-R18M-K19. RNA was diluted to microinject concentrations of 200-1000 pg in 5 nl pulses for the following constructs: pCS2-mCherry-R18, pCS2-mCherry-R18M, pCS2-C-cadherin-eGFP, pcDNA3.1-FLAG-HA-14-3-3-β, and pCS2-mem-RFP. Embryos utilized for microscopy were injected dorsally into both blastomeres at two-cell stage to target mesendoderm. Embryos utilized for immunoprecipitation were injected at one-cell stage in the animal cap.

### Immunoprecipitation

Whole embryos or dissected mesendoderm for immunoprecipitation were processed in aforementioned lysis buffer with 1% Tergitol type NP-40 (endogenous IP) or 1% Triton X-100 (FLAG IP). Endogenous IP lysates (equivalent to 40 embryos) were initially incubated with 50 μl Protein-G agarose bead slurry (Roche Diagnostics, 11243233001) for an hour. Beads were pelleted by centrifugation at 3000 rpm for 5 minutes (4°C). The IP lysate was then removed and incubated with primary antibody (5 μg Pan 14-3-3, Santa Cruz) for overnight immunoprecipitation. The Protein-G beads were washed three times in lysis buffer for ten minutes. Bead samples were stored in 2x Laemmli buffer with 5% β-mercaptoethanol at −80°C for subsequent gel electrophoresis. After overnight primary incubation, endogenous IP lysates were incubated with 50 μl Protein-G agarose bead slurry overnight. FLAG IP lysates (equivalent to 30 embryos) were incubated with 40 μl agarose slurry covalently linked to anti-FLAG M2 mAb (Sigma-Aldrich, A2220) for overnight immunoprecipitation. IP samples were pelleted by centrifugation at 3000 rpm for 5 minutes (4°C), and lysates were removed for storage as supernatant samples. Beads were washed with three exchanges of lysis buffer (1% Tergitol type NP-40 for endogenous IP beads; 1% Triton X-100 for FLAG IP beads) for ten minutes. IP bead samples were stored in 25 μl of 2x Laemmli buffer with 5% β-mercaptoethanol, incubated on a 95°C heat block for five minutes, and loaded onto 12% SDS-PAGE polyacrylamide gels for electrophoresis. Gels were either stained using Sypro Red or transferred onto nitrocellulose membrane prior to incubation with antibodies. All incubation and wash steps were performed at 4°C using a vertical rotator. All centrifugation steps were performed at 4°C.

### LC/MS-MS Proteomic Analysis

Prior to staining with Sypro Red dye (Invitrogen, S12000) SDS-PAGE gels were incubated twice in fixative solution consisting of 50% methanol and 7% glacial acetic acid for 30 minutes per incubation. After decanting the second fixative solution, the gel was placed in a fresh dish and incubated in 60 ml of Sypro Red dye overnight. The staining solution was decanted and the gel was incubated in a wash solution of 10% methanol and 7% glacial acetic acid for 30 minutes. Afterwards, the gel was washed three times in 100 ml of commercial ultrapure water before gel imaging and further preparation for LC/MS-MS. All incubations and washes were performed at room temperature on a flat rotator.

LC/MS-MS was performed by the Center for Advanced Proteomics Research (CAPR) at the Rutgers New Jersey Medical School. Sypro Red-labeled SDS-PAGE gel sections were excised at the facility and in-gel trypsin digestion was performed. The resulting peptides were C18 desalted and analyzed by LC/MS-MS on the Q Exactive instrument. The MS/MS spectra were searched against the NCBI *Xenopus laevis* database using MASCOT (v.2.3) search engines on the Proteome Discoverer (V1.4) platform. The protein false discovery rate is less than 1%. The mass spectrometry data were obtained from an Orbitrap instrument funded in part by NIH grant NS046593, for the support of the UMDNJ Neuroproteomics Core Facility. Information in tables was derived utilizing Scaffold 4.7.3 with a protein and peptide false discovery rate of 1%.

### Mesendoderm Dissection and Dissociation

Stage 10.5 embryos in 0.5x MBS were dissected at the animal cap surface to reveal the mesendodermal mantle. The dorsal side of the embryo was identified and the mesendoderm leading edge was excised from the rest of the tissue. These cells were pipette transferred into a solution of Ca^2+^/Mg^2+^ free 1xMBS on 1% Ca^2+^/Mg^2+^ free agarose and allowed to dissociate for 30 minutes at room temperature. Dissociated cells were transferred to fibronectin (Sigma-Aldrich, F4759) matrix (200 μl of 1:7.5 in water; incubated on MatTek glass bottom microwell dishes, P35G-1.5-14-C) and imaged. Fibronectin incubation was performed overnight at 4°C.

### Dorsal Marginal Zone Explant Preparation

DMZ explants were excised and plated as described in (Davidson *et al.*, 2004). Stage 10.5 embryos in 0.5x MBS were carefully dissected at the animal cap surface to reveal the mesendodermal mantle. The embryo was then bisected and the dorsal side of the embryo was used. The remaining animal cap cells and endodermal cells were trimmed away, leaving the mesoderm, mesendoderm, and bottle cells of the embryo. The explant was flattened on fibronectin (Sigma-Aldrich, F4759) matrix (100 μl of 1:7.5 in water; incubated on MatTek glass bottom microwell dishes, P35G-1.5-14-C) and silicone grease was used to mount a coverslip. Fibronectin incubation was performed overnight at 4°C. Explants were slightly compressed and allowed to attach and migrate for 2 hours (live imaging experiments) or 5 hours (samples to be fixed). In order to fix explants, the coverslip was removed just prior to incubation in 100% methanol.

### Immunofluorescence

Embryos and dorsal marginal zone explants were fixed in ice-cold 100% methanol or Dent’s fixative (80% methanol, 20% DMSO) and incubated at −20°C overnight. Embryos were rehydrated in partial changes of 0.1x Modified Barth’s Saline (MBS). Embryos were transferred into vinyl molds with dimensions of 15 mm × 15 mm × 5 mm (Electron Microscopy Sciences, 62534-15) containing tissue freezing medium (Electron Microscopy Sciences, 72592), and briefly immersed in liquid nitrogen to flash freeze. Frozen embryo blocks were stored at −80°C until sectioned (Leica 819, 14035838925; 40 μm per slice) at −20°C using a temperature controlled cryostat and mounted on slides (VWR, 48311-703). Prior to blocking, embryo sections and explants were rehydrated in 1x TBS. Samples were permeablized in 1x TBS with 0.25% Triton X-100 for 10 minutes before blocking in 5% goat serum for one hour at room temperature. Incubation in primary antibody (30 minutes at 37°C) was followed by three buffer exchanges with 1x TBS and incubation in Alexa Fluor 488 or 555 conjugated goat anti-mouse (Invitrogen, A11029; A21424) and/or goat anti-rabbit (Invitrogen, A11034; A21429) IgG (30 minutes at 37°C). Explants were imaged after three buffer exchanges with 1x TBS. Embryonic sections were serially dehydrated using 1x TBS with increasing percentages of methanol and cleared using benzyl benzoate/benzyl alcohol. Slides were coverslipped (Corning, 2980-245), sealed with multiple coats of nail polish, and left for drying overnight (4°C) before imaging.

### Linescan Analysis and Quantification

Linescan measurements of fluorescence were conducted using the linescan tool in the profile tab of the Zen 2.3 lite software application. The first measurement was taken from the approximate center of a cell and extended to the area proximal to C-cadherin signal (eGFP). The second measurement was taken from the approximate center of a cell and extended into the C-cadherin signal. The first and last five measurements in the line scan (a length of approximately 0.5 μm) were averaged and a ratio of the average at the end of the line to the average at the beginning of the line was calculated for each cell. Four ratios were determined for each cell; signal proximal to cell-cell adhesion/center (eGFP and RFP), and signal at cell-cell adhesion/center (eGFP and RFP). To determine if a statistical difference in the means of these two ratios was present, a paired samples t-test was conducted.

### FRAP Experiments

Filaments chosen for photobleaching were imaged every fifteen seconds over the course of one minute to capture intensity of fluorescence prior to bleaching. The filament was then bleached using 100% power on the 488 channel for a duration of two minutes. The area of photobleaching was determined by a capture of the photobleaching mask acquired at the end of the two minute bleach, along with the capture of the filament segment with reduced fluorescence at the start of the recovery period. Images of recovery of fluorescence within the bleach zone were captured in ten second intervals beginning after the conclusion of the bleach period.

Recovery of fluorescence was measured using the Zen 2.3 lite application. A region of interest (ROI) was aligned to the bleached segment of the filament in each of the post-bleach time points to determine the mean value of fluorescent intensity (mFI). This ROI was also used to measure the intensity of the bleach zone before photobleaching. An ROI was similarly used to measure a non-bleached control area of the filament proximal to the bleach zone. Using the method described by (Zheng *et al.*, 2011) with minor alterations, the rate of photobleaching (rate r) was first established by determining the ratio of the control ROI mFI in each post-bleach time point to the average mFI across pre-bleach control ROIs [*r* = *mFI*_*c*_ ÷ *mFI*_*c*0_]. The normalized fluorescent intensity (NFI) was established by determining the ratio of the difference between the mFI of the initial experimental post-bleach time point and each subsequent experimental post-bleach time point to the rate established for that time point [*NFI* = (*mFI*_*b*#_ − *mFI*_*b*1_) ÷ *r*]. The recovery percentage (rec) of each time point was established by determining the ratio of the NFI in each experimental post-bleach time point to the average mFI across experimental prebleach ROIs and multiplying that ratio by 100 [*rec*_#_ = *NFI*_#_ ÷ (*mFI*_b_0) × 100]. Time point series to be measured for recovery of fluorescence in this way were first scrutinized for clarity of measurement. Exclusion criteria for recovery of fluorescence experiments included unclear bleach zones, measurement zones that were obstructed during the series by crossing or merging filaments, gross reorganization of filaments that resulted in loss of the measurement zone, and readily apparent movement of the measurement zone and proximal regions such that they did not remain in the plane of focus.

Analysis of fluorescent recovery was conducted by plotting the percentage of recovery for each post-bleach time point for a given series against time increments (seconds) and fitting a trendline to the data. Variation in time was controlled for by using the same time duration for each analysis (20 seconds to 330 seconds). The slope of the trendline was determined for each analysis and the mean slope was compared across R18 and R18M groups using a one-way t-test in which the sample variances were shown to be equal via an F-test.

### Gap quantification

Z-stacks of eGFP-K19 expressing mesendoderm cells after collision were observed for the presence of a filament depleted zone proximal to a cell-cell contact. These gaps were counted and a proportion of total gaps to total cells was calculated for cells expressing mCh-R18 and cells expressing mCh-R18M. In order to detect if a statistical difference in these ratios was present, a z-test for proportions from two samples was utilized. The length of gaps from the adhesion to the first visible filaments was measured using the linescan tool in the profile tab of the Zen 2.3 lite software application.

### Image Acquisition

Images for live and fixed samples were taken using a Zeiss Observer spinning disk confocal microscope using a 40x 1.3 NA or 63x 1.4 NA Apochromat objective, unless specified otherwise.

## Acknowledgements

We thank the members of the Weber Laboratory and the Rutgers Department of Biological Sciences for their useful feedback during the course of this work. We also thank the many investigators who provided reagents for these studies. We thank Drs. Tong Liu and Hong Li of the New Jersey Medical School Center for Advanced Proteomics Research for their assistance in processing and obtaining the mass spectrometry data. Mass spectrometry data were obtained from an Orbitrap instrument funded in part by NIH grant NS046593, for the support of the UMDNJ Neuroproteomics Core Facility. This research was supported by USPHS grant R15-HD084254 to G.F.W.

## Supplemental Material

**Supplemental Figure 1.**
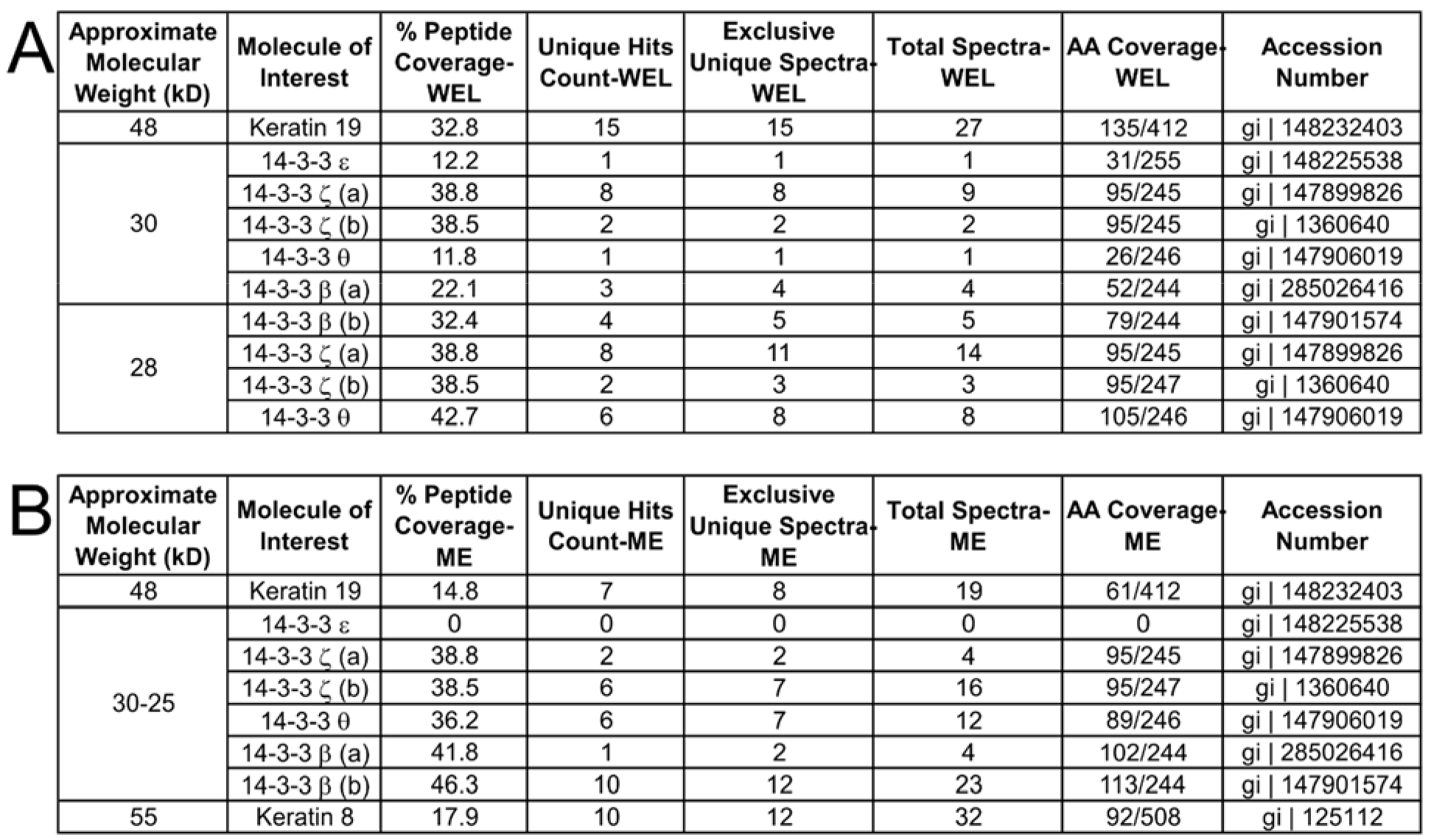
Extended proteomic analysis. A-B) Tables of selected proteomic information from LC/MS-MS analysis of 14-3-3 immunoprecipitated samples (Fig. 2). Table A includes data from whole embryo lysate samples. Table B includes data from mesendoderm only samples. Analysis was performed using Scaffold 4.7.3.

**Supplemental Figure 2.**
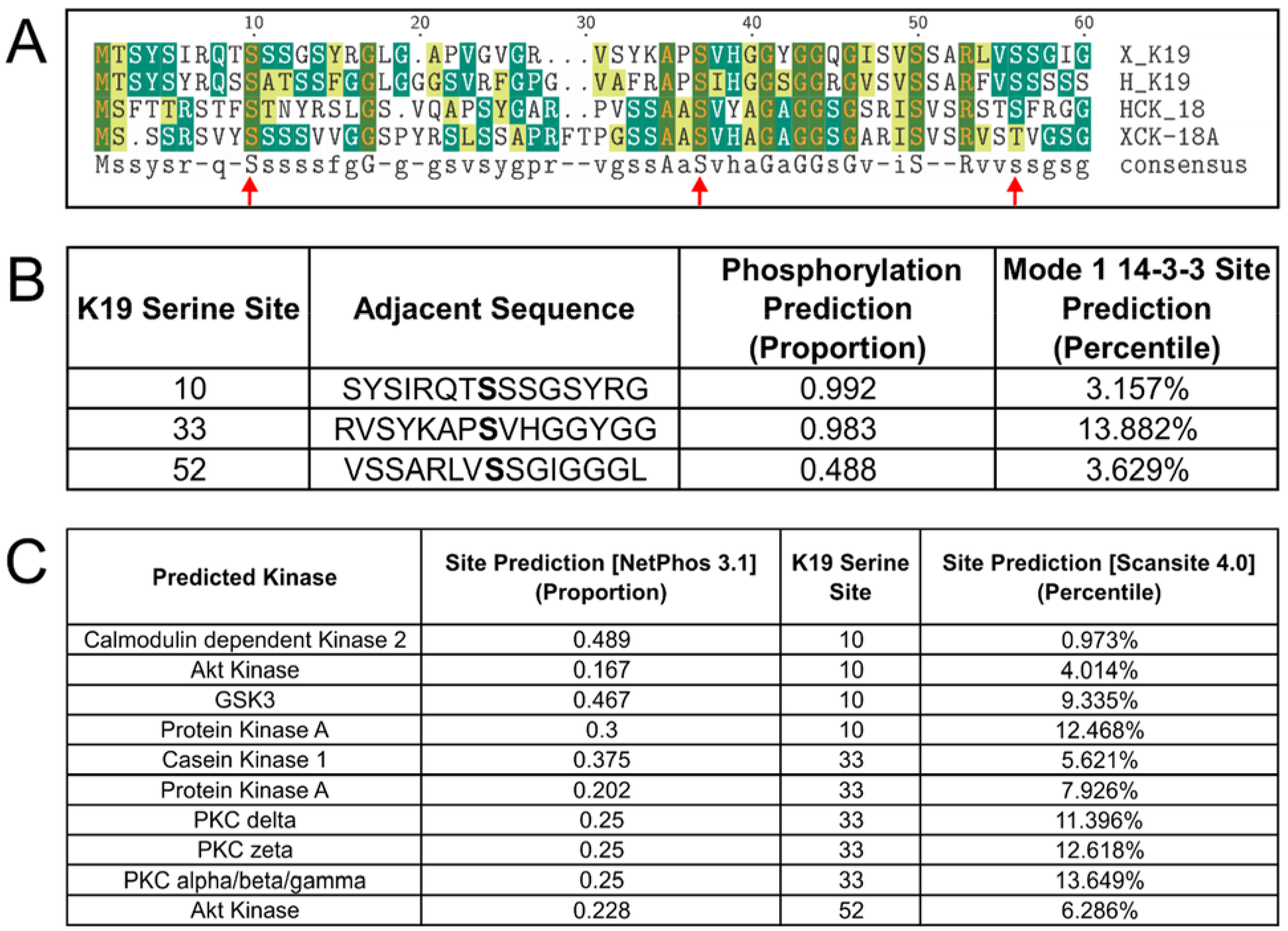
14-3-3 informatics. A) Sequence alignment of *Xenopus laevis* Keratin 18A and 19 with human Keratin 18 and 19. The arrows indicate serine sites referenced in B and C. The sequence alignment was created using the San Diego Supercomputer (SDSC) Biology Workbench. B) Table reporting the likelihood that a given site is phosphorylated (NetPhos 3.1) (Blom *et al.*., 1999) and is a 14-3-3 site (Scansite 4.0) (Obenauer *et al.*., 2003). C) Table reporting the likelihood that a given site is a kinase site. Reported results were restricted to sites identified in both NetPhos3.1 (Blom *et al.*., 2004) and Scansite 4.0 analyses. NetPhos 3.1 proportions display the likelihood that a given site is a true site relative to how close the score is to 1. Lower Scansite 4.0 percentiles (closer to 0%) represent greater likelihood that a given site is a true site. NetPhos 3.1 likelihoods in A are for unspecified kinases. Scansite 4.0 was utilized at the minimum stringency to report all predictions at the selected sites.

**Supplemental Figure 3.**
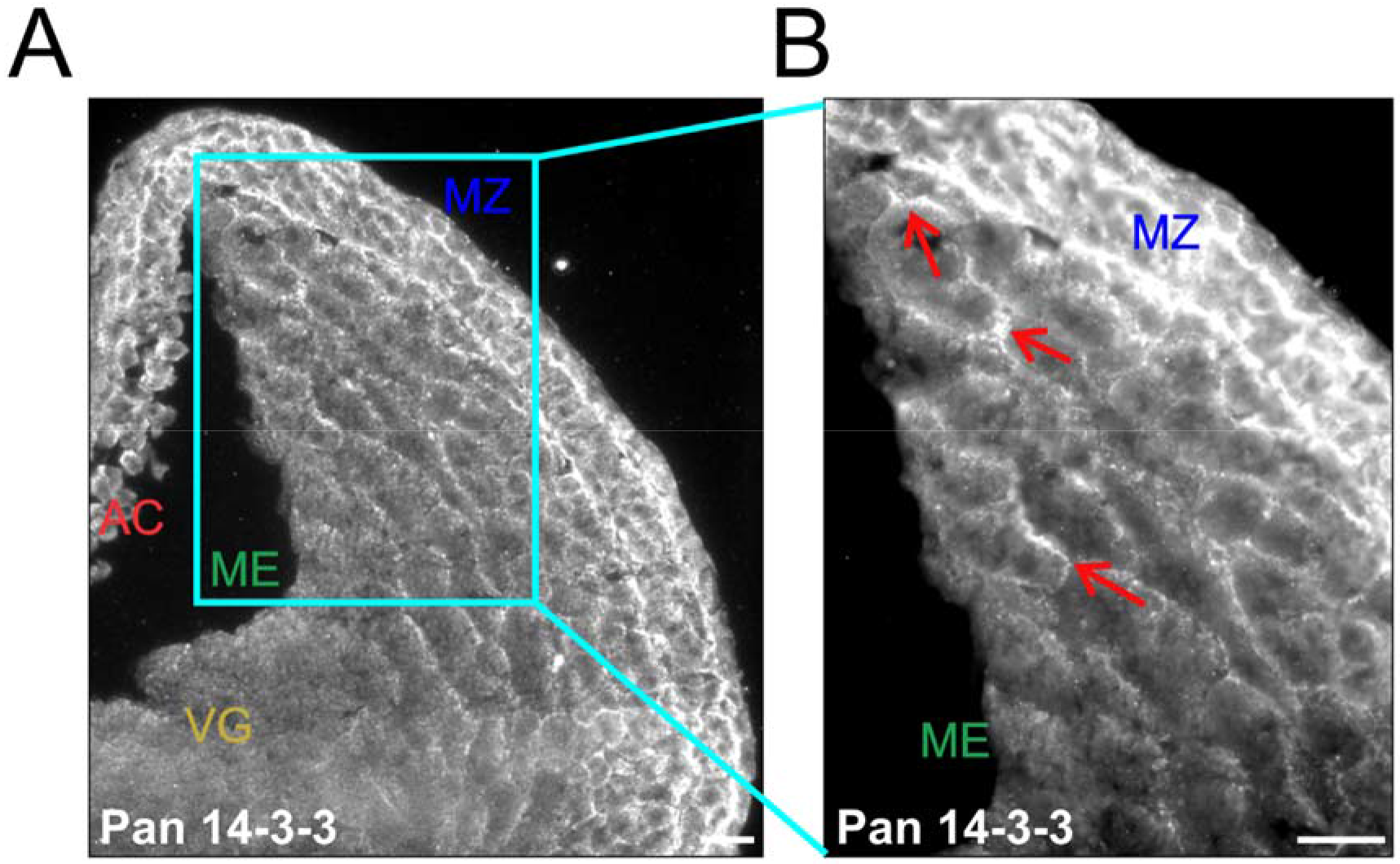
Low magnification of 14-3-3 immunofluorescence. A) Whole *Xenopus* embryos were cryosectioned at stage 10.5 (40 μm thick slices) and immunolabeled using an antibody that detects multiple 14-3-3 isoforms. Labels refer to the animal cap (AC), mesendoderm (ME), marginal zone (MZ), and vegetal hemisphere (VG) of the embryo. Image taken using 10x objective (0.25 NA). B) Region labeled by the cyan box in panel A in increased magnification. Arrows indicate rear contacts of cells where 14-3-3 labeling is prominent. Image taken using 20x objective (0.75 NA). Scale bars are 50 μm.

**Supplemental Movie 1.** FRAP movies. Recovery of eGFP-K19 fluorescence was measured for mesendoderm explants expressing mCherry-R18M (Movie A) and mCherry-R18 (Movie B). Photobleaching was performed using the 488 laser at 100% power in the area indicated by the ROI box for 120 seconds. Images were taken every 10 seconds after photobleaching (330 seconds total). Scale bars are 10 μm.

## <u>References</u>

Acehan, D., Petzold, C., Gumper, I.,Sabatini, D. D., Mueller, E. J., Cowin, P., and Stokes, D. L. (2008). Plakoglobin Is Required for Effective Intermediate Filament Anchorage to Desmosomes. J. Invest. Dermatol. 128, 2665–2675.

Boudreau, A., Tanner, K., Wang, D., Geyer, F. C., Reis-Filho, J. S., and Bissell, M. J. (2013. 14-3-3sigma stabilizes a complex of soluble actin and intermediate filament to enable breast tumor invasion. Proc Natl Acad Sci U S A 110, E3937–44.

Cheung, K. J., Gabrielson, E., Werb, Z., and Ewald, A. J. (2013). Collective invasion in breast cancer requires a conserved basal epithelial program. Cell. 155, 1639–1651.

Colakoğlu, G., and Brown, A. (2009). Intermediate filaments exchange subunits along their length and elongate by end-to-end annealing. J. Cell. Biol. 185, 769–777.

Conway, D. E., Breckenridge, M. T., Hinde, E., Gratton, E., Chen, C. S., and Schwartz, M. A. (2013). Fluid shear stress on endothelial cells modulates mechanical tension across VE-cadherin and PECAM-1. Curr. Biol. 23, 1024–1030.

Davidson, L. A., Keller, R., and DeSimone, D. (2004). Patterning and tissue movements in a novel explant preparation of the marginal zone of Xenopus laevis. Gene Expr. Patterns 4, 457466.

Deng, M. et al. (2013). Lactotransferrin acts as a tumor suppressor in nasopharyngeal carcinoma by repressing AKT through multiple mechanisms. Oncogene. 32, 4273–4283.

le Duc, Q., Shi, Q., Blonk, I., Sonnenberg, A., Wang, N., Leckband, D., and de Rooij, J. (2010). Vinculin potentiates E-cadherin mechanosensing and is recruited to actin-anchored sites within adherens junctions in a myosin II-dependent manner. J. Cell. Biol. 189, 1107–1115.

Fradette, J., Germain, L., Seshaiah, P., and Coulombe, P. A. (1998). The type I keratin 19 possesses distinct and context-dependent assembly properties. J. Biol. Chem. 273, 35176–35184.

Heasman, J., Ginsberg, D., Geiger, B., Goldstone, K., Pratt, T., Yoshida-Noro, C., and Wylie, C. (1994). A functional test for maternally inherited cadherin in Xenopus shows its importance in cell adhesion at the blastula stage. Development. 120, 49–57.

Herrmann, H., Häner, M., Brettel, M., Müller, S. A., Goldie, K. N., Fedtke, B., Lustig, A., Franke, W. W., and Aebi, U. (1996). Structure and assembly properties of the intermediate filament protein vimentin: the role of its head, rod and tail domains. J. Mol. Biol. 264, 933–953.

Hofmann, I., and Franke, W. W. (1997). Heterotypic interactions and filament assembly of type I and type II cytokeratins in vitro: viscometry and determinations of relative affinities. Eur. J. Cell Biol. 72, 122–132.

Jin, J. et al. (2004). Proteomic, functional, and domain-based analysis of in vivo 14-3-3 binding proteins involved in cytoskeletal regulation and cellular organization. Curr. Biol. 14, 1436–1450.

Johnson, C., Crowther, S., Stafford, M. J., Campbell, D. G., Toth, R., and MacKintosh, C. (2010). Bioinformatic and experimental survey of 14-3-3-binding sites. Biochem. J. 427, 69–78.

Ju, J. H. et al. (2015). Cytokeratin19 induced by HER2ERK binds and stabilizes HER2 on cell membranes. Cell Death Differ. 22, 665–676.

Kim, S., Wong, P., and Coulombe, P. A. (2006). A keratin cytoskeletal protein regulates protein synthesis and epithelial cell growth. Nature. 441, 362–365.

Kirmse, R., Portet, S., Mücke, N., Aebi, U., Herrmann, H., and Langowski, J. (2007). A quantitative kinetic model for the in vitro assembly of intermediate filaments from tetrameric vimentin. J. Biol. Chem. 282, 18563–18572.

Klymkowsky, M. W., Shook, D. R., and Maynell, L. a 1992. Evidence that the deep keratin filament systems of the Xenopus embryo act to ensure normal gastrulation. Proc. Natl. Acad. Sci. U. S. A. 89, 8736–8740.

Kolsch, A., Windoffer, R., Wurflinger, T., Aach, T., and Leube, R. E. (2010). The keratin-filament cycle of assembly and disassembly. J. Cell. Sci. 123, 2266–2272.

Ku, N. O., Liao, J., and Omary, M. B. (1998). Phosphorylation of human keratin 18 serine 33 regulates binding to 14-3-3 proteins. Embo J. 17, 1892–1906.

Lau, J. M. C., Wu, C., and Muslin, A. J. (2006). Differential role of 14-3-3 family members in Xenopus development. Dev. Dyn. 235, 1761–1776.

Lee, C.-H., Kim, M.-S., Chung, B. M., Leahy, D. J., and Coulombe, P. A. (2012). Structural basis for heteromeric assembly and perinuclear organization of keratin filaments. Nat. Struct. Mol. Biol. 19, 707–715.

Li, H. H., Guo, Y., Teng, J. L., Ding, M. X., Yu, A. C. H., and Chen, J. G. (2006). 14-3-3 gamma affects dynamics and integrity of glial filaments by binding to phosphorylated GFAP. J. Cell. Sci. 119, 4452–4461.

Liao, J., and Omary, M. B. (1996). 14-3-3 proteins associate with phosphorylated simple epithelial keratins during cell cycle progression and act as a solubility cofactor. J. Cell. Biol. 133, 345–357.

Loschke, F., Homberg, M., and Magin, T. M. (2016). Keratin Isotypes Control Desmosome Stability and Dynamics through PKC&alpha; J. Invest. Dermatol. 136, 202–213.

Loschke, F., Seltmann, K., Bouameur, J. E., and Magin, T. M. (2015). Regulation of keratin network organization. Curr. Opin. Cell. Biol. 32, 56–64.

Magin, T. M., Schröder, R., Leitgeb, S., Wanninger, F., Zatloukal, K., Grund, C., and Melton, D. W. (1998). Lessons from keratin 18 knockout mice: Formation of novel keratin filaments, secondary loss of keratin 7 and accumulation of liver-specific keratin 8-positive aggregates. J. Cell. Biol. 140, 1441–1451.

Margolis, S. S. et al. (2006). Role for the PP2A/B565 Phosphatase in Regulating 14-3-3 Release from Cdc25 to Control Mitosis. Cell. 127, 759–773.

Maruthamuthu, V., Sabass, B., Schwarz, U. S., and Gardel, M. L. (2011. Cell-ECM traction force modulates endogenous tension at cell-cell contacts. Proc. Natl. Acad. Sci. U. S. A. 108, 4708–4713.

Megason, S. G., and Fraser, S. E. (2003). Digitizing life at the level of the cell: high-performance laser-scanning microscopy and image analysis for in toto imaging of development. Mech. Dev. 120, 1407–1420.

Mertz, A. F., Che, Y., Banerjee, S., Goldstein, J. M., Rosowski, K. A., Revilla, S. F., Niessen, C. M., Marchetti, M. C., Dufresne, E. R., and Horsley, V. 2013. Cadherin-based intercellular adhesions organize epithelial cell-matrix traction forces. Proc. Natl. Acad. Sci. U. S. A. 110, 842–847.

Miao, L. Q., Teng, J. L., Lin, J. Q., Liao, X. Z., and Chen, J. G. (2013). 14-3-3 proteins interact with neurofilament protein-L and regulate dynamic assembly of neurofilaments. J. Cell. Sci. 126, 427–436.

Muslin, A. J., Tanner, J. W., Allen, P. M., and Shaw, A. S. (1996). Interaction of 14-3-3 with signaling proteins is mediated by the recognition of phosphoserine. Cell. 84, 889–897.

Nelson, C. M., and Chen, C. S. (2003). VE-cadherin simultaneously stimulates and inhibits cell proliferation by altering cytoskeletal structure and tension. J. Cell. Sci. 116, 3571–3581.

Nieuwkoop, P. D. (Pieter D.., and Faber, J. (1994). Normal table of Xenopus laevis (Daudin): a systematical and chronological survey of the development from the fertilized egg till the end of metamorphosis, Garland Pub.

Nöding, B., Herrmann, H., and Köster, S. (2014). Direct Observation of Subunit Exchange along Mature Vimentin Intermediate Filaments. Biophys. J. 107, 2923–2931.

Obsil, T., and Obsilova, V. (2011). Structural basis of 14-3-3 protein functions. Semin. Cell Dev. Biol. 22, 663–672.

Omary, M. B., Ku, N. O., Tao, G. Z., Toivola, D. M., and Liao, J. (2006). “Heads and tails” of intermediate filament phosphorylation: multiple sites and functional insights. Trends Biochem. Sci. 31, 383–394.

Petosa, C., Masters, S. C., Bankston, L. A., Pohl, J., Wang, B. C., Fu, H. I., and Liddington, R. C. (1998). 14-3-3 zeta binds a phosphorylated Raf peptide and an unphosphorylated peptide via its conserved amphipathic groove. J. Biol. Chem. 273, 16305–16310.

Ridge, K. M., Linz, L., Flitney, F. W., Kuczmarski, E. R., Chou, Y. H., Omary, M. B., Sznajder, J. I., and Goldman, R. D. (2005). Keratin 8 phosphorylation by protein kinase C delta regulates shear stress-mediated disassembly of keratin intermediate filaments in alveolar epithelial cells. J. Biol. Chem. 280, 30400–30405.

Riveline, D., Zamir, E., Balaban, N. Q., Schwarz, U. S., Ishizaki, T., Narumiya, S., Kam, Z., Geiger, B., and Bershadsky, A. D. (2001). Focal contacts as mechanosensors: Externally applied local mechanical force induces growth of focal contacts by an mDia1-dependent and ROCK-independent mechanism. J. Cell. Biol. 153, 1175–1185.

Roberts, B. J., Reddy, R., and Wahl, J. K. (2013). Stratifin (14-3-3 sigma) Limits Plakophilin-3 Exchange with the Desmosomal Plaque. PLoS One 8, 14.

Sanghvi-Shah, R., and Weber, G. F. (2017). Intermediate filaments at the junction of mechanotransduction, migration, and development. Front. Cell Dev. Biol. 5.

Sehgal, L. et al. (2014). 14-3-3 gamma-mediated transport of plakoglobin to the cell border is required for the initiation of desmosome assembly in vitro and in vivo. J. Cell. Sci. 127, 2174–2188.

Sivaramakrishnan, S., Schneider, J. L., Sitikov, A., Goldman, R. D., and Ridge, K. M. (2009). Shear Stress Induced Reorganization of the Keratin Intermediate Filament Network Requires Phosphorylation by Protein Kinase C zeta. Mol. Biol. Cell 20, 2755–2765.

Snider, N. T., and Omary, M. B. (2014). Post-translational modifications of intermediate filament proteins: mechanisms and functions. Nat. Rev. Mol. Cell Biol. 15, 163–177.

Sonavane, P. R., Wang, C., Dzamba, B., Weber, G. F., Periasamy, A., and DeSimone, D. W. (2017). Mechanical and signaling roles for keratin intermediate filaments in the assembly and morphogenesis of Xenopus mesendoderm tissue at gastrulation. Development. 144, 4363–4376.

Strnad, P., Windoffer, R., and Leube, R. E. (2002). Induction of rapid and reversible cytokeratin filament network remodeling by inhibition of tyrosine phosphatases. J. Cell. Sci. 115, 4133–4148.

Torpey, N., Wylie, C. C., and Heasman, J. (1992). Function of maternal cytokeratin in Xenopus development. Nature. 357, 413–415.

Tzivion, G., Luo, Z. J., and Avruch, J. (2000). Calyculin A-induced vimentin phosphorylation sequesters 14-3-3 and displaces other 14-3-3 partners in vivo. J. Biol. Chem. 275, 29772–29778.

Vikstrom, K. L., Lim, S. S., Goldman, R. D., and Borisy, G. G. (1992). Steady-state dynamics of intermediate filament networks. J. Cell. Biol. 118, 121–129.

Vishal, S. S., Tilwani, S., and Dalal, S. N. (2018). Plakoglobin localization to the cell border restores desmosome function in cells lacking 14-3-3γ. Biochem. Biophys. Res. Commun. 495, 1998–2003.

Wang, B., Yang, H., Liu, Y. C., Jelinek, T., Zhang, L., Ruoslahti, E., and Fu, H. (1999). Isolation of high-affinity peptide antagonists of 14-3-3 proteins by phage display. Biochemistry. 38, 12499–12504.

Weber, G. F., Bjerke, M. A., and DeSimone, D. W. (2012). A Mechanoresponsive Cadherin-Keratin Complex Directs Polarized Protrusive Behavior and Collective Cell Migration. Dev. Cell 22, 104–115.

Windoffer, R., Beil, M., Magin, T. M., and Leube, R. E. (2011). Cytoskeleton in motion: the dynamics of keratin intermediate filaments in epithelia. J. Cell. Biol. 194, 669–678.

Windoffer, R., Woll, S., Strnad, P., and Leube, R. E. (2004). Identification of novel principles of keratin filament network turnover in living cells. Mol. Biol. Cell 15, 2436–2448.

Winklbauer, R., Selchow, A., Nagel, M., and Angres, B. (1992). Cell interaction and its role in mesoderm cell migration during Xenopus gastrulation. Dev. Dyn. 195, 290–302.

Woll, S., Windoffer, R., and Leube, R. E. (2007). p38 MAPK-dependent shaping of the keratin cytoskeleton in cultured cells. J. Cell. Biol. 177, 795–807.

Yaffe, M. B., Rittinger, K., Volinia, S., Caron, P. R., Aitken, A., Leffers, H., Gamblin, S. J., Smerdon, S. J., and Cantley, L. C. (1997). The structural basis for 14-3-3: phosphopeptide binding specificity. Cell. 91, 961–971.

Zheng, C.-Y., Petralia, R. S., Wang, Y.-X., and Kachar, B. (2011). Fluorescence Recovery After Photobleaching (FRAP) of Fluorescence Tagged Proteins in Dendritic Spines of Cultured Hippocampal Neurons. JOVE-JOURNAL Vis. Exp.

Zhou, Q. Q. et al. (2010). 14-3-3 Coordinates Microtubules, Rac, and Myosin II to Control Cell Mechanics and Cytokinesis. Curr. Biol. 20, 1881–1889.

Zhou, X., Liao, J., Hu, L., Feng, L., and Omary, M. B. (1999). Characterization of the major physiologic phosphorylation site of human keratin 19 and its role in filament organization. J Biol Chem 274, 12861–12866.

## <u>References</u>

Blom, N., Gammeltoft, S., and Brunak, S. (1999). Sequence and structure-based prediction of eukaryotic protein phosphorylation sites. J. Mol. Biol. 294, 1351–1362.

Blom, N., Sicheritz-Pontén, T., Gupta, R., Gammeltoft, S., and Brunak, S. (2004). Prediction of post-translational glycosylation and phosphorylation of proteins from the amino acid sequence. Proteomics. 4, 1633–1649.

Obenauer, J. C., Cantley, L. C., and Yaffe, M. B. (2003). Scansite 2.0: proteome-wide prediction of cell signaling interactions using short sequence motifs. Nucleic Acids Res. 31, 3635–3641.

